# Non-standard proteins in the lens of AlphaFold 3 - a case study of amyloids

**DOI:** 10.1101/2024.07.09.602655

**Authors:** Alicja W. Wojciechowska, Jakub W. Wojciechowski, Gert Vriend, Malgorzata Kotulska

## Abstract

While three-dimensional structures of globular and transmembrane forms are available for many amyloid proteins, structures of their amyloid forms are scarce in the PDB. Amyloids pose major challenges for both experimental structure determination and computational modelling.

We evaluated the amyloid-modelling performance of the current top modelling software, AlphaFold 3 (AF3), using three datasets. Dataset 1 contains 153 proteins and peptides that are known to form fibrils, but their 3D structures have not been experimentally determined. Dataset 2 contains 56 non-aggregating/non-amyloid peptides. Dataset 3 contains seven proteins for which the three-dimensional fibrillar structure is known.

Fibrillar structures were predicted for 34% of dataset 1, but unfortunately also for 54% of dataset 2. Fibrillar structures were successfully predicted for five out of seven proteins from dataset 3. Comparing AF3 with different methods, it outperformed Boltz, and predicted the structures of CsgA and α-synuclein more correctly than RibbonFold, whereas the latter predicted Aβ-42 better.

The performance of AF3 in prediction of amyloid structures for our datasets seems hindered by low abundance of amyloid structures in the PDB and high prevalence of structure data for their non-fibrillar forms. AF3 tends to assign a higher quality score to globular oligomeric models than to fibrillar ones. A correct amyloid structure prediction is more likely to be obtained for shorter fragments. The amyloid modelling quality of AF3 seems underwhelming, but it can still provide hypotheses about amyloid structures in some cases. Our work also suggests the steps needed to achieve a better performance in the near future.

**Statement for a broader audience:** Amyloid proteins can form stable, insoluble fibrils that are often related to a neurodegenerative disease. Knowledge of the three-dimensional structure of these fibrils is important, e.g. for a drug design. We evaluate the performance of AlphaFold 3 on the prediction of amyloid structures and observe that it struggles with these cases. The problems seem to arise mainly from the nature of the AlphaFold 3 training dataset and polymorphic nature of many amyloids. Although the results are underwhelming, AlphaFold 3 can sometimes provide valuable insights into amyloid protein structures, something that only a few years ago still seemed a very hard to reach goal.

## Introduction

Parkinson’s disease, Alzheimer’s disease, and several more diseases, which have a big influence on our society, are associated with proteins whose mutations enhance the predisposition for a disease. Good three-dimensional models of their structures can help understand the disease states (Efraimidis et al. 2023). Some of these proteins can exist in two forms (Volles and Lansbury 2003; Soto 2003, (Jiang and Eisenberg 2025). One form is a nonfibrillar one, which can be globular, transmembrane, or disordered. The other form is an amyloid fibril that in many cases tends to be cytotoxic. A prominent example of an amyloid protein is α-synuclein, which natively exists in either a disordered state or in a helical form when it binds to lipid. In the amyloid form it builds Lewy body inclusions in Parkinson’s disease patients (Bartels, Choi, and Selkoe 2011).

Not all amyloids are disease-related. Many organisms produce amyloid fibrils that have physiological functions. For example, amyloid fibrils of curli (CsgA) protein serve as scaffolds for biofilm matrices of enteric bacteria (Otzen and Riek 2019), filamentous fungi use the amyloid fold of Het-s to control immune responses (Daskalov et al. 2012; Daskalov 2016), and amyloid fibrils of human Pmel17 regulate the pigmentation in melanocytes (Watt et al. 2013).

Information about the three-dimensional structure of amyloid proteins can shed light on the pathology of amyloid-related diseases and can support drug discovery (Iadanza et al. 2018; Lai et al. 2023). Amyloids are hard to address with biophysical techniques. Kinetics studies require that sample preparation follows complex dissolution protocols that often do not lead to perfect monomerization (Faller and Hureau, 2020). Amyloids are highly sensitive to environmental conditions such as pH and temperature, so that full experimental control over their aggregation is difficult (Lutter, Aubrey, and Xue 2021). Structure determination with NMR or X-ray is troublesome due to the big size of amyloid fibrils and the possible presence of multiple polymorphic species in the sample (Zielinski, Röder, and Schröder 2021; Toyama and Weissman 2011). Cryo Electron Microscopy shows great promise for amyloid structure determination, but has not yet led to a flurry of amyloid structures (Creekmore, Chang, and Lee 2021). Consequently, the representation of amyloids in the PDB (Protein Data Bank) (Burley et al. 2017) is poor, except for a few well-known examples like Aβ-42 (and Aβ-40) or α-synuclein.

Amyloid proteins often display polymorphism. Fragments of α-synuclein and Aβ-42, for example, can form nanotubular cross-beta architectures or tape-like structures (Morris et al. 2013; Lu et al. 2003), and IAPP (amylin) even changes its fibrillar structure several times during amyloid assembly (Wilkinson et al. 2023).

The majority of software in the amyloid field focuses on the identification of aggregation-prone regions (APRs) (for reviews, see e.g. (Kotulska and Wojciechowski 2022; Iglesias et al. 2024). WALTZ predicts APRs using position-specific scoring matrices (Maurer-Stroh et al. 2010). TANGO is rooted in statistical mechanics (Fernandez-Escamilla et al. 2004). FoldAmyloid calculates the probability of hydrogen-bonding and expected packing density of the residues (Garbuzynskiy, Lobanov, and Galzitskaya 2010). PATH combines physicochemical information with machine learning (Wojciechowski and Kotulska 2020). Amylogram uses n-gram analysis (Burdukiewicz et al. 2017), and Aggrescan 3D relies on structural information to find surface-accessible APRs (Kuriata et al. 2019). Cross-Beta is a very recent amyloidogenic protein predictor based on machine learning that uses different sizes of the sliding window (Gonay et al. 2025).

Few tools attempt to address the problem of modelling 3D structures of amyloids. Fibpredictor predicts the structure of amyloid fibrils by estimating the energy scores of the known fibril architectures of a protein (Tabatabaei Ghomi, Topp, and Lill 2016). BetaSerpentine gives guidance on how beta-arches are preferentially placed in the fibril structure (Tabatabaei Ghomi, Topp, and Lill 2016; Bondarev et al. 2018). The authors of RibbonFold trained an AlphaFold-like neural network to predict an amyloid fibril structure for any protein sequence (Guo et al. 2025).

The release of AlphaFold 2 (AF2) by DeepMind stunned the modelling world in CASP 14, where it performed so well that many insiders in the field called the protein structure prediction problem ‘solved’ (Subramaniam and Kleywegt 2022; Thornton, Laskowski, and Borkakoti 2021), at least for single-domain proteins. The article describing AF2 has already been cited >35.000 times (Jumper et al. 2021) and AF3 more than 8000 times (Abramson *et al*. 2024). Neither AF2 nor AF3 require the structure of a homologous protein to be known in order to build a model, which could make it the perfect tool for modelling amyloid proteins. For this task, AF3 seems to be a more appropriate choice than AF2 due to lower reliance on MSA, and generative character that could help to predict multiple structures for amyloid proteins.

Amyloid proteins has been envisioned as a significant challenge for AF2 due to their polymorphic nature (Pinheiro *et al*. 2021). We confirm the same for AF3, although to a lesser extent, showing that it also struggles with the prediction of amyloid fibrils. Our results suggest that the root of the problem lies in the small number of amyloid structures in the protein database (PDB), combined with the fact that for many proteins that can form an amyloid, a non-fibrillar conformation (of a homolog) exists, and it could have been used for AF3 training.

## Results

### Choosing oligomeric state (number of protein copies, n)

AF3, the most recent version of AF, can predict protein structures including interactions between molecular entities (Jumper et al. 2021; Abramson et al. 2024). This is important as the presence or absence of other entities can influence the protein’s conformation, like active versus inactive form (Bywater 2013). The presence of either GDP or GTP, for example, determines if the inactive or active form of a G protein should be modelled, while the presence of calcium for many enzymes determines that they should be modelled in the active form, *et cetera*. The switch between the monomeric form (e.g. the globular native state) and the amyloid form of a protein tends to be much more complicated than the mere absence or presence of an ion or a cofactor.

The oligomeric state input parameter (number of protein copies, *n*) is the main parameter to inform AF3 that an amyloid fibril structure should be modelled. As it is computationally infeasible to always run the AF3 server using a large number of different values for *n*, we tested the influence of this input parameter on the quality of an amyloid structure models. The tests were performed on the Aβ-42 peptide, to find one good-for-all value for *n*. Aβ-42 was chosen because it is well studied, and many PDB files are available for both monomers and fibrils. It should be noted that polymorphisms are observed for the monomer and for the fibrils of Aβ-42 (Roche et al. 2016; Barrow et al. 1992; Zhang et al. 2000).

Fifty Aβ-42 models were generated for each value of *n* from 1 through 9. Monomers (*n*=1) invariably were predicted as helical. For dimers (*n*=2), helices, sheets, and mixed structures were predicted. All models for *n*>2 had fibril-like architectures (see Figure 1A).

**Figure 1.**
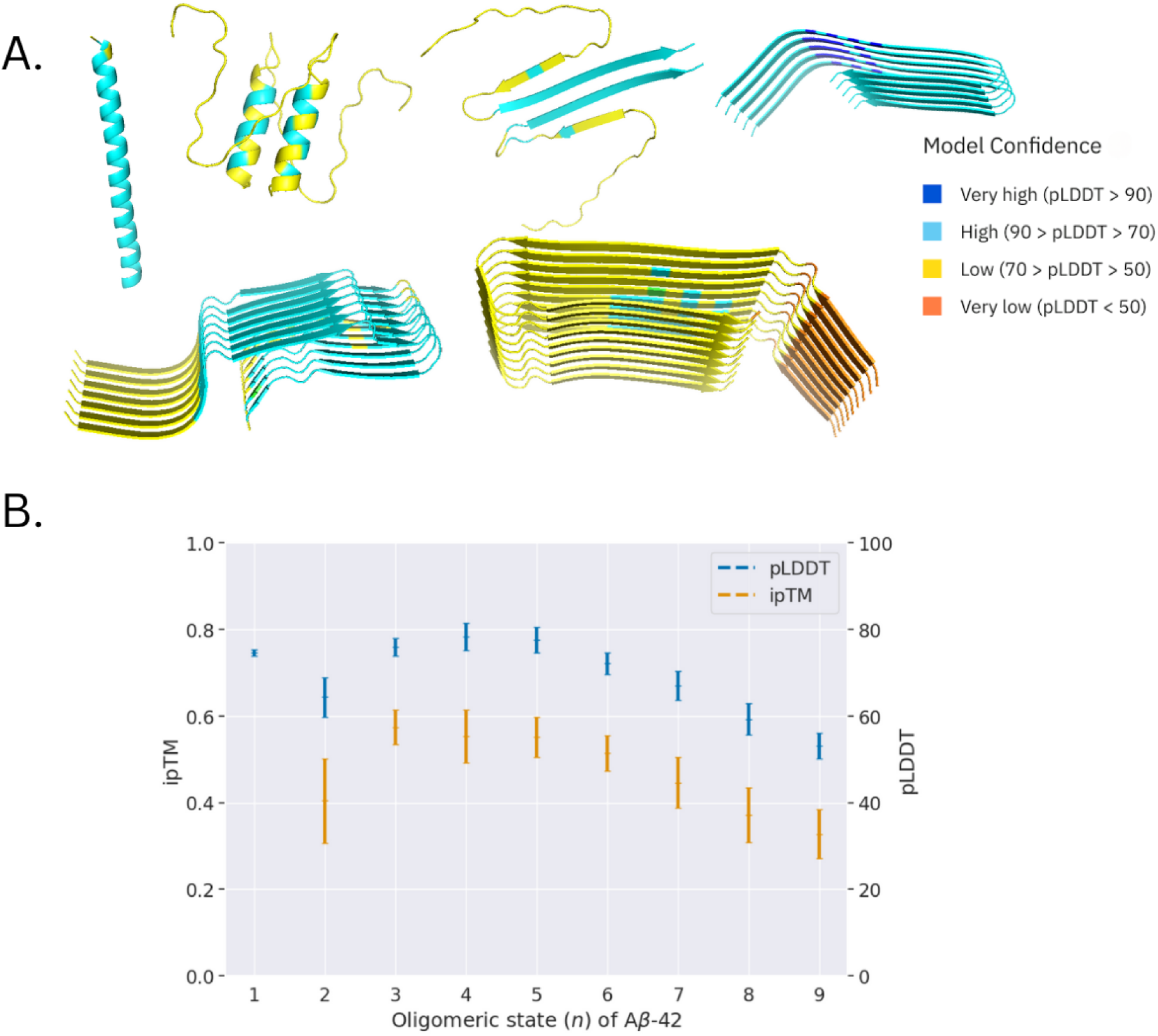
Impact of oligomeric state, n, on Aβ-42 structure prediction with AF3. **A.** Typical Aβ-42 models colored by pLDDT metric. From left to right and top to bottom: monomer, helical dimer, beta-sheet dimer, and fibrillar structures for n=5, n=7, and n=9, respectively. **B.** pLDDT (blue) and iPTM (orange) as a function of n value.

AF3, as well as AF2, provide several measures related to the predicted quality of models. The pLDDT relates to how well residues are modelled locally, according to the AF internal evaluation, and the ipTM is related to the predicted quality of multimeric interfaces. The Aβ-42 models for *n*=1 can be considered confident (mean pLDTT = 75). The highest pLDDT and ipTM values were obtained for *n*=4 and *n*=5 (see Figure 1B). It should be noted that experimental scientists tend to deposit amyloid fibrils as repeats of typically 6 or more copies, which is thus also what AF3 obtained for its training (see Figure S1). We therefore chose to predict all amyloid models, in the remainder of this work, with oligomeric state *n*=6 as a compromise between the predicted model quality and the typical oligomeric state observed in PDB files.

### AF3 struggles to predict fibrillar structures

AF3 was tested on 153 proteins known to be amyloid but without known experimental structure (*Positive-control* dataset) and on 56 peptides known to be non-amyloid (*Negative-control* dataset). Our *Negative-control* dataset represents a group of sequences that are often used to train amyloid sequence predictors.

No full 3D structures of the 153 amyloid fibrils are known, but what is known, though, is that they can form fibrils. Therefore, a prediction is called *correct* if any amyloid structure is predicted in the model, while every other prediction is called *wrong*. Conversely, predictions for the 56 non-amyloid peptides are called *wrong* if any amyloid fibril is predicted for them by AF3, and otherwise they are called *correct*. The pLDDT and ipTM distributions of models for the *Positive-* and *Negative-control* datasets are presented in Figures S2B and S2D. Models with C6 symmetry, helical structures, and complex non-amyloid multimers were obtained. Figure 2A shows some typical examples. In Figure S3 we present an example where the predictions went completely *wrong* for three different peptides derived from one protein. Figure 2B shows the confusion matrix for the predictions. 54% of the peptides known to be non-aggregating were *wrongly* modelled as fibrils, while only 34% of the proteins and peptides known to be aggregating were *correctly* predicted as such.

**Figure 2.**
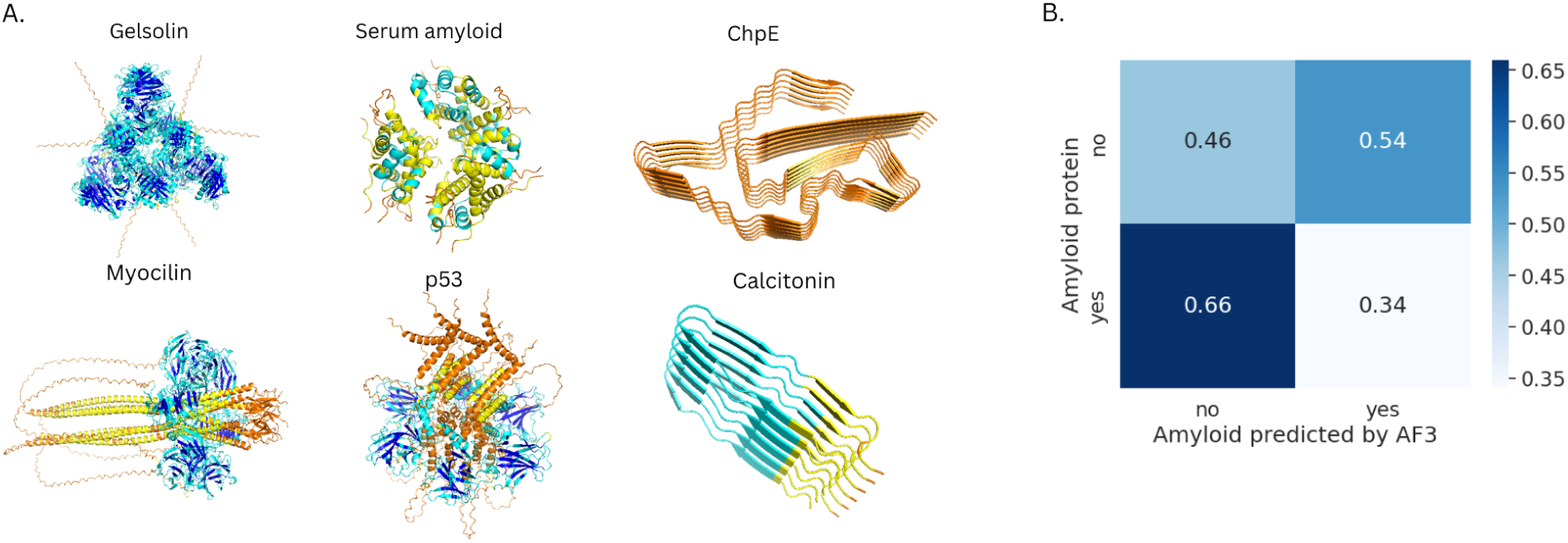
Amyloid structure prediction results. **A.** A few typical results. Top: Gelsolin model with C6 symmetry, helical model of Serum amyloid, and amyloid fibrillar model of ChpE. Bottom: non-amyloid model of Myocilin, non-amyloid model of p53, and amyloid fibril model of Calcitonin. **B.** Confusion matrix for the 209 predictions for Positive-control and Negative-control.

*Correctly* predicted sequences from the *Positive-control* dataset were, on average, two and a half times shorter than those *wrongly* predicted (an average length of 70 and 189 residues for the *correct* and *wrong* predictions, respectively; this difference is statistically significant: p-value=6e-6 for the Mann-Whitney test (see also Figures S4 and S5). The pLDDT of *wrong* predictions was on average 18 points higher than the pLDDT of *correct* predictions (Mann-Whitney test, p-value=1e-5). No statistically significant difference was observed for ipTM. These results suggest that AF3 tends to have more faith in its own models when they are globular than when they are amyloid. Notably, the longer the sequence, the smaller the chance that AF3 will model it as an amyloid fibril.

When AF3 predicts a protein structure, by default it will search for a possible template in the PDB to guide the prediction. When we switched this flag off in AF3, three more sequences (out of 153) in the *Positive-control* dataset were *correctly* predicted as fibrils.

### Monomers versus multimers

Does AF model a multimer by combining individually modelled monomers, or does it model a multimer as a single entity? The latter seems preferred for modelling amyloid fibrils. To verify this, a monomer from each protein in the *Positive-control* dataset was modelled, and compared with one unit from the corresponding model of *n*=6. Our analyses showed that 56 out of the 153 monomers were similar, with a TM-score>0.5, to at least one unit in the corresponding multimer according to Foldseek. TM-score, provided by Foldseek, is a measure of the structural similarity between protein structures (structures are considered significantly similar when their TM-score is > 0.5 (Xu and Zhang 2010)) (van Kempen et al. 2024). Figure 3 shows the confusion matrix for the predictions for the *Positive-control* dataset, illustrating the relationship between the monomer and the most similar unit from the multimer (*n*=6). High similarity between the monomeric and multimeric models was associated with a low chance for *correct* amyloid structure prediction. Only 13% of the *correct* multimeric models shared similarities with their corresponding monomer. In contrast, 51% of the *wrong* multimeric models were similar to their corresponding monomer.

**Figure 3.**
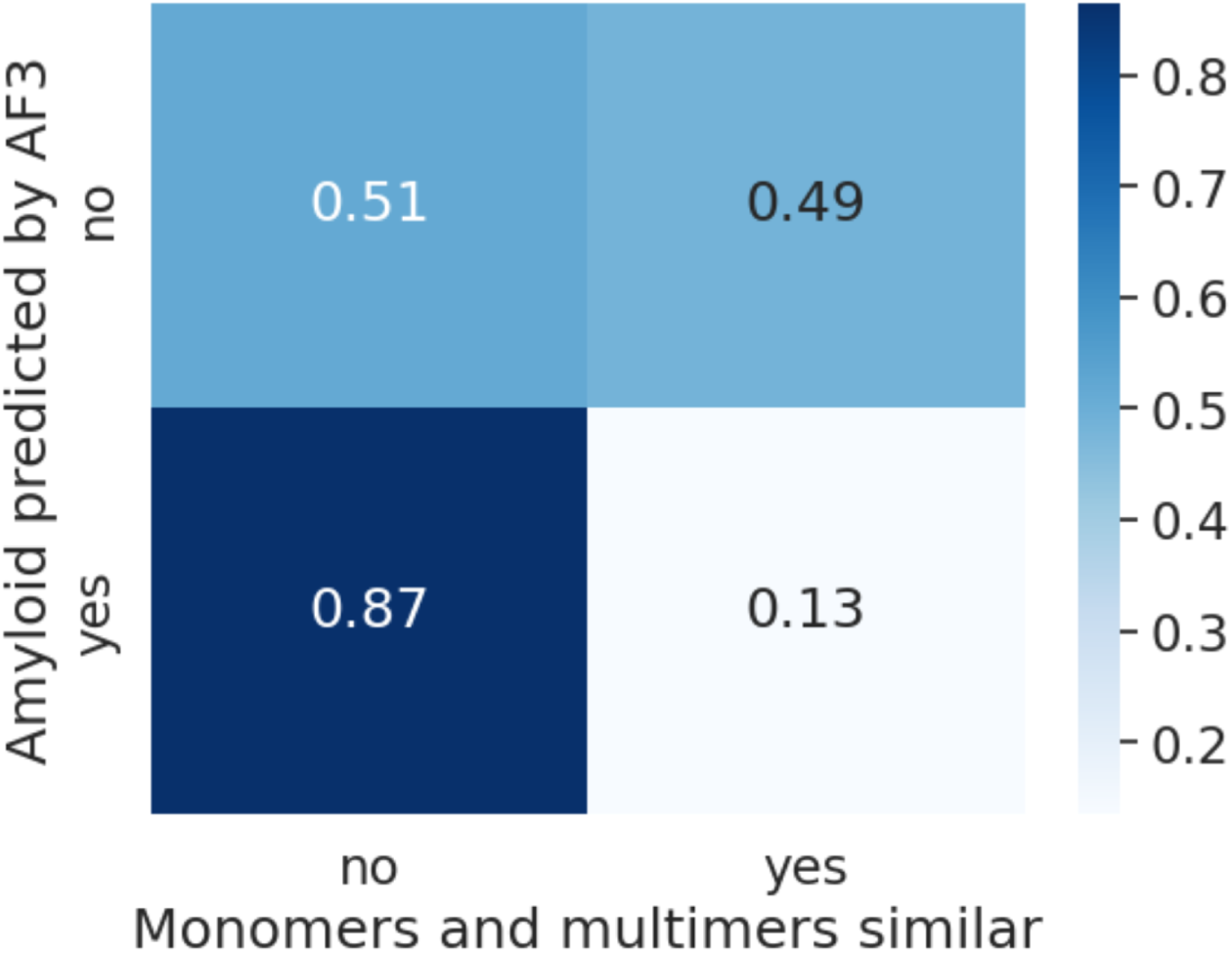
Confusion matrix for the 153 amyloid proteins predictions (Positive-control dataset) and similarity (TM-score>0.5) between monomers and multimers.

High similarity between the monomeric and multimeric form was also associated with a considerably higher pLDDT (15 points higher on average; Mann-Whitney, p-value=9e-10). These effects were stronger for longer proteins (see Figure S6), which is in line with the aforementioned sequence length effects.

### Bias caused by homologs in the PDB

The training data of AF2 and AF3 consisted mainly of globular proteins and contained only very few amyloids. This might be one of the reasons why AF3 struggles with amyloids. For each *n*=6 model from the *Positive-control* dataset, the most similar structure in the PDB was identified with Foldseek (details on the Foldseek runs are given in the Methods section).

In 72 out of the 101 *wrong* models, predictions resembled at least one PDB structure with TM-score>0.5. A majority of the other 29 models were too short to allow for the detection of a significant hit. Most of the PDB hits were found as similar to globular homologs of the *Positive-control* proteins. 75% of these hits had a significant Foldseek homolog probability (above 0.8). (According to Foldseek’s documentation, the homology probability is the “estimate of the probability for a query and target to be homologous, e.g. being within the same SCOPe superfamily”). This effect was stronger for longer sequences (Mann-Whitney, p-value=2e-12, see Figure S7) and associated with higher pLDDT scores (Mann-Whitney, value=0.01). A few representative examples of cases where AF3 *wrongly* predicted a globular model are presented in Figure 4.

**Figure 4.**
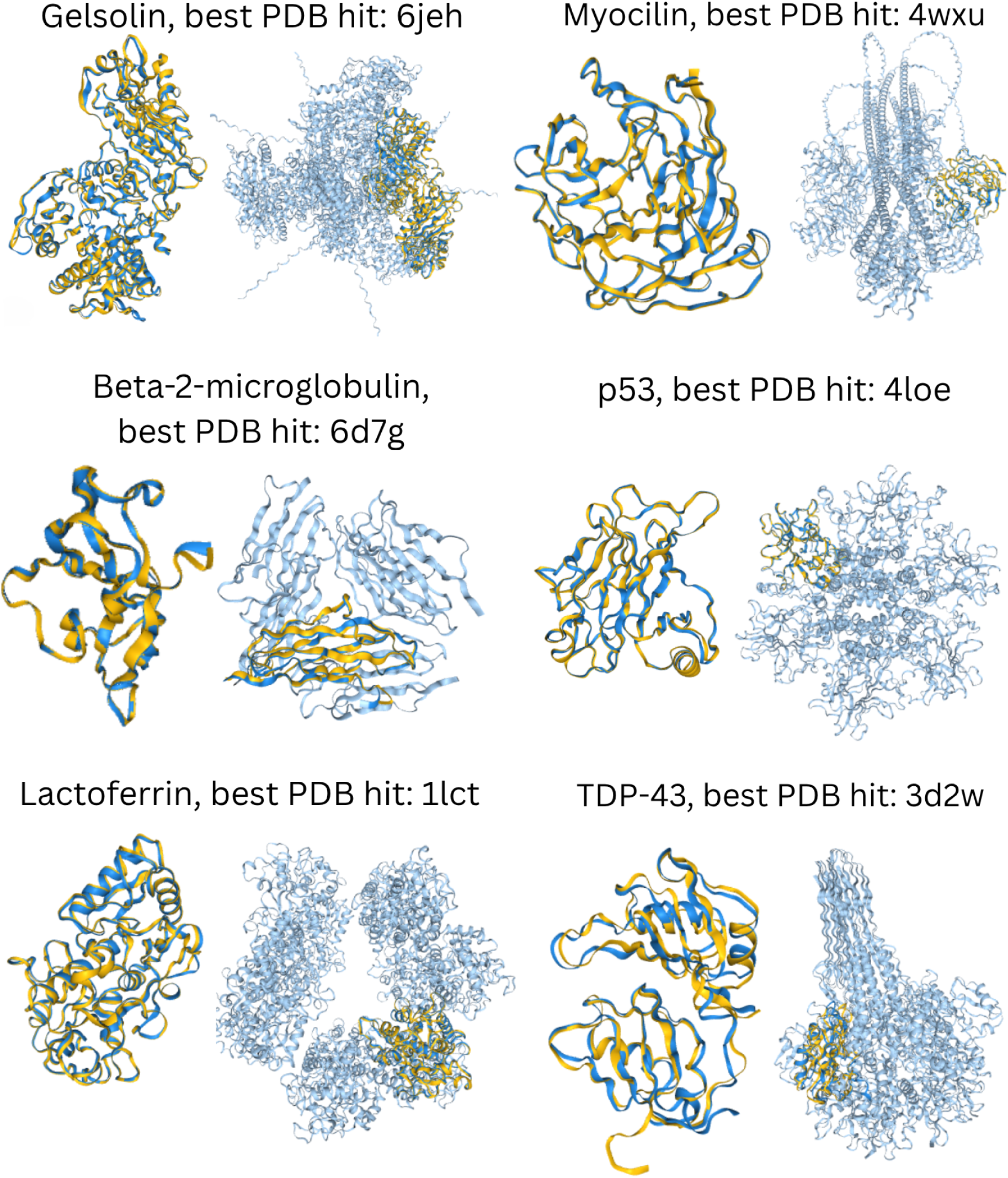
Six representative examples of structure alignments between wrong Positive-control models (blue) and their most similar PDB file - the one with the highest TM-score (yellow) for: Gelsolin (TM-score=0.99), Myocilin (TM-score=1.0), Beta-2-microglobulin (TM-score=1.0), p53 (TM-score=1.0), Lactoferrin (TM-score=1.0), TDP-43 (TM-score=1.0). The right-hand panel shows the superposition results, while the left-hand panel shows the superposed parts only. The Foldseek homolog probability was 1.0 for all examples.

Foldseek found hits in the PDB for 10 out of 52 *correct* predictions for the *Positive-control* data (see Table 1). Two of these ten *correct* models were related to homologs in the PDB with a significant Foldseek homolog probability: CsgB and protegrin. The model of CsgB was similar to its ortholog CsgA. However, the CsgA structure was not seen by AF3 (structure released after AF3 ‘cutoff date’); therefore, modelling of CsgB could have not been biased by this example. The PDB structure of protegrin had some amyloid characteristics (β-sheets) that could have guided AF3 toward an amyloid structure. No other *correct* prediction was significantly similar to a fibrillar structure that could have been seen by AF3 during training. Therefore, 51 out of the 52 *correct* predictions seem to have been *de novo*.

**Table 1.** Foldseek hits found for the correct Positive-control models in the PDB.

| Protein | Monomer and multimer similar? [tmscore] | PDB identifier of the hit | TM-score for one chain from the n=6 model and the PDB hit | Foldseek homolog probability | Comment about the hit PDB file |
| --- | --- | --- | --- | --- | --- |
| CsgB | yes [0.84] | 8enr | 0.9 | 1.0 | solved amyloid fibril of the ortholog (not seen by AF3) |
| Protegrin | yes [0.6] | 2mz6 | 0.6 | 0.7 | solved dimer that has beta sheets |
| PL B1 protein (95-114aa) | no [0.4] | 2jor | 0.66 | 0.3 | shared subsequence, monomer with beta sheets |
| MspA | yes [0.9] | 1mkj | 0.6 | 0.02 | not homolog, pdb has some beta sheets |
| Spectrin (68-47aa) | yes [0.7] | 5bnd | 0.8 | 0.3 | not homolog, pdb has some beta sheets |
| Acyl phosphatase (34-53aa) | yes [0.55] | 4p24 | 0.7 | 0.3 | not homolog, pdb contains beta barrel |
| FapB | yes [0.8] | 8enr | 0.2 | 0.02 | not homolog, amyloid fibril |
| FapC | yes [0.84] | 5awf | 0.25 | 0.01 | not homolog, complex with beta sheets |
| ODAM | no [0.1] | 4l06 | 0.4 | 0.04 | not homolog, pdb has some beta sheets |
| ChpE | no [0.3] | 8a00 | 0.25 | 0.01 | not homolog, amyloid fibril |

MMseqs2 (Mirdita, Steinegger, and Söding 2019) was used to search for homologs in the PDB for the sequences of *correct* models. Nine hits were found (see Table S1) to a protein with a globular structure. For these nine cases, AF3 not only *correctly* predicted an amyloid structure, but it even overcame a globular signal from its training data. Table S1 includes a summary for these nine cases.

Generally, AF3 struggles with the bias in its training data which misleads its predictions of amyloid proteins (Figure 4), though in some cases we observed that it can go beyond PDB memorization (Table 1 and S1).

### Predicting amyloids for which the structure is already known

Can AF3 improve on amyloid structure predictions if we add (more) amyloid structure data to its training set? We tried to address this question by predicting 50 monomeric (*n*=1) and 50 multimeric (*n*=6) models for each of the six amyloid fibril structures in the PDB release that was used for AF3 training: Aβ-42, α-synuclein, glucagon, IAPP (human amylin), transthyretin, and immunoglobulin. The pLDDT and ipTM scores are shown in Figure S2A and S2B, respectively. AF3 predicted structures with pLDDT scores that indicated *low* to *confident* quality. Table 2 gives a summary of the results.

**Table 2.**
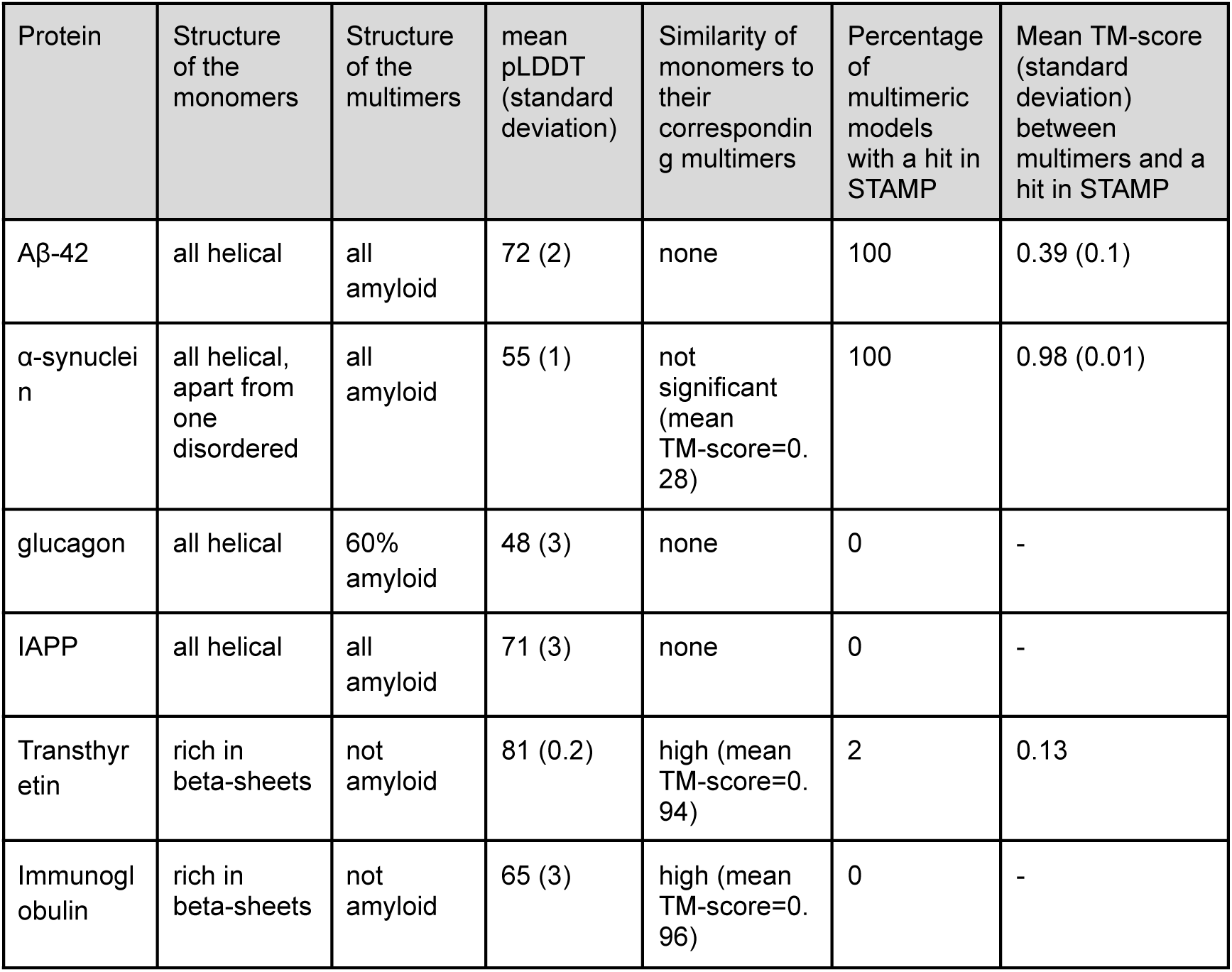
Summary of the results of AF3 predictions for the ResolvedAmyloidStructure dataset. AF3 models were generated for monomeric (n=1) and multimeric (n=6) forms of each protein and compared to each other. Multimeric models (n=6) were compared with the amyloid structures found in STAMP for the corresponding protein using Foldseek.

Foldseek found two clusters in the monomer structure ensemble of IAPP and just one in the other five ensembles. The monomer models of Aβ-42, α-synuclein, glucagon, and IAPP were all helical, apart from one disordered structure of α-synuclein. This is in line with the experimental data on the structures of these proteins.

Foldseek found one cluster in the multimeric ensemble of Aβ-42, IAPP, immunoglobulin, and transthyretin; it found two clusters for α-synuclein and eight for glucagon. The cluster representatives are shown in Figure S8. Visual inspection of the multimer models revealed amyloid structures for Aβ-42, α-synuclein, and IAPP. A fibrillar pattern could be found in 60% of the glucagon models. The models of immunoglobulin did not resemble an amyloid, and neither did those of transthyretin.

We compared monomeric structures to their multimeric counterparts for the proteins in the *ResolvedAmyloidStructure* dataset. For each representative structure from the 50 monomeric models, the multimer containing the most similar constituent monomers was identified. Monomeric models of Aβ-42, IAPP, glucagon, and α-synuclein were not similar to any of their multimeric counterparts, whereas the monomeric structures of immunoglobulin and transthyretin were highly similar to the multimeric counterparts. We compared the AF3 models with their corresponding amyloid structures (see Figure S9) based on STAMP database of amyloid structures (Louros et al. 2022). Only the hits regarding the same protein were considered. Hits with a TM-score>0.5 could only be found for models of Aβ-42 and α-synuclein.

We studied in detail the two cases for which the least accurate modelling performance was observed: immunoglobulin and transthyretin, and using Foldseek scanned the whole PDB for structures matching their multimeric models. For each model, a non-fibrillar match with TM-score>0.9 was found. This suggests that globular structures of these proteins could have biased AF3 towards non-fibrillar models (see list of hits in Figure S10). Additionally, we tested whether the MSA choice (default AF3 or ColabFold generated) and its alignment depth would make a difference (see Table S3). All models from the AF3 web server, all desktop generated AF3 models with default MSA, and all desktop AF3 models with MSAs of different depth could still be clustered altogether into one group with a nonfibrillar representative (see Figure S11).

When AF3 has multiple examples of monomeric and multimeric forms of amyloid proteins in its training set, it can distinguish between them, and the models of α-synuclein, Aβ-42, IAPP, and glucagon were different for monomers and multimers. Nevertheless, the similarity of models to their corresponding amyloid structures was generally low, except for all models of α-synuclein and 5 models of Aβ-42. Although the IAPP models had a β-hairpin architecture typical of amyloids, these did not match the real fibrillar structures of this protein (see Figure S8 and S9). We speculate that the poor performance for transthyretin and immunoglobulin results from a low number of amyloid structures and a very large number of globular structures in the AF3 training set.

We separately studied the CsgA protein. The CsgA amyloid structures, 8enr and 8enq, were deposited in the PDB after the cutoff date of AF3 training. Note that in PDB, we can find structures of very short fragments of CsgA, including amyloid structures deposited before AF3 cutoff date. The largest fragment solved of CsgA and seen by AF3 has 22 amino acids and is part of 6l7c.

The mean pLDDT of the 50 CsgA multimeric models with *n=*6 was 78 (Figure S8), and all models could be grouped into one cluster (Figure S8). For each multimeric model of CsgA, we found a match to the 8enr structure (see Figure S9D) with a TM-score greater than 0.99. No structure matched 6l7c. In short, AF3 modelled the full-sequence-length CsgA amyloid structure correctly.

### Modeling software other than AF3 web server

We compared the performance of AF3 web server to its desktop version, and to other methods predicting protein structure, by applying them to the *ResolvedAmyloidStructure* dataset and to CsgA.

For the *ResolvedAmyloidStructure* dataset and CsgA, structures with an oligomeric state of six (*n*=6), were modeled with ColabFold using the third version of AF-Multimer (Mirdita et al. 2022). We analysed the performance of AF-Multimer with three protocols: 1) using an automatically chosen template from PDB100; 2) using an amyloid structure as a custom template; 3) no template was used. Chains in the models were not aligned in a perpendicular manner typical of amyloids in none of the cases (see Figure S12-S14). As the poor performance of AF-Multimer on amyloid proteins has also been observed by (Ragonis-Bachar, 2024), we did not follow up with an extensive analysis for these models.

We generated ten models, using different software for each protein in the *ResolvedAmyloidStructure* dataset and for CsgA, following four protocols:

1. AF3 desktop version with *n*=6;
2. AF3 desktop version with custom MSA (obtained from ColabFold) and *n*=6;
3. Boltz with *n*=6 (Passaro et al. 2025; Wohlwend et al. 2025);
4. RibbonFold (*n*=5) (Guo et al. 2025).

Unless mentioned otherwise, the default parameters were used. We compared all models to their corresponding amyloid structures in STAMP (see Figure 5). For CsgA, their 8enr and 8enq structures were added to the analysis.

**Figure 5.**
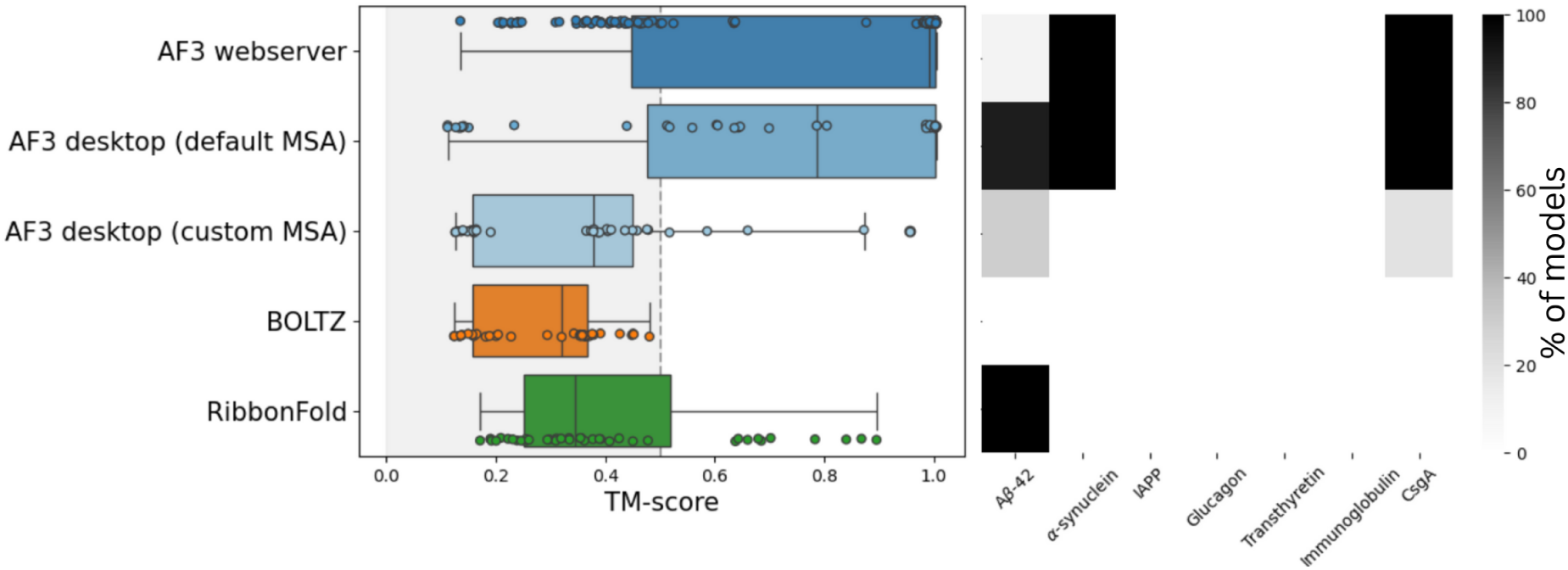
Comparison of the performance of structure prediction algorithms. Models were aligned with Foldseek to the corresponding amyloid structures in STAMP. Left: Boxplots of TM-score values between the model and the best hit. Right: percentage of models (grey colour intensity) with a TM-score>0.5 for each protein in the ResolvedAmyloidStructure dataset and CsgA for each protocol is given.

Not one Boltz model had a TM-score>0.5 for any of the corresponding real amyloid structure. RibbonFold outperformed AF3 in the case of Aβ-42, with the mean TM-score between the model and actual Aβ structures equal to 0.74 (standard deviation = 0.1) (see also Figure S15). Only AF3 managed to predict the fibrillar structure of α-synuclein and CsgA with a TM-score>0.95. The web server version and the AF3 desktop version with default MSA usually provided similar results, though they differed slightly for Aβ-42, for which the desktop version performed a little better. The AF3 desktop version with custom MSA gave the worst results of all AF3 protocols.

## Discussion

Amyloid proteins are often characterised by structural polymorphism, differences between monomeric and multimeric forms, low solubility, and sensitivity of their aggregation propensity to environmental conditions, post-translational modifications and interactions. Consequently, the structures of amyloids are hard to address, both experimentally and *in silico*. AF3 is the first software that allows for unsupervised modelling of amyloids, including and their monomeric and multimeric forms. Unfortunately, modelling of amyloid structures still poses a challenge to all methods, including Alpha Fold. Amyloid models are of much lower quality than the results of modelling non-amyloid proteins by AF2 and AF3’s. Only 34% of known amyloids were predicted with AF3 as such, while 56% of known non-amyloids were falsely predicted as amyloid fibrils. Upon comparison of the two versions, we observed that AF3 is to be preferred over AF2 for amyloid structure prediction, as the latter failed to generate fibrillar structures for our tested sequences (*ResolvedAmyloidStructure* dataset) (see also (Ragonis-Bachar et al. 2024).

Evidently, AF3 heavily relies on the data seen during itstraining and this appears to be the main reason why AF3 struggles with proteins that can adopt several different conformations when data is only available for one of those. This problem has been observed previously for both AF2 and AF3 (Chakravarty et al. 2024; Chakravarty et al. 2025; Chakravarty and Porter 2022). We demonstrated that this effect is especially strong for amyloids, as their structures are severely underrepresented in the PDB. Many amyloids of our study were predicted as globular proteins that resembled their homologs seen by AF3 during its training. Another underlying cause of the poor modelling accuracy could be that amyloid sequence–structure relationships may not conform very well to Anfinsen’s thermodynamic hypothesis, as indicated by the pronounced polymorphism of many amyloids and their very high sensitivity to environmental conditions (Jiang and Eisenberg 2025).

Incorrect AF3 (and AF2) models, attributed with user-misleading high predicted model qualities (pLDDT scores), have already been observed for protein dimers (Schmid and Walter 2025), fold-switching proteins (Schafer and Porter 2025), and beta-solenoids (Pratt et al. 2025). We observed a strong association between the model pLDDT and TM-score to its most similar PDB entry. This result explains why incorrect globular predictions for amyloid proteins tend to have a higher pLDDT value than those of correct fibrillar models. Metrics other than pLDDT are needed to assess the predicted quality of amyloid structure models, but perhaps for predictions of other proteins too.

Encouragingly, however, in our study in a few cases AF3 demonstrated an ability to generalize beyond ‘memorising’ the (entire) PDB. This potential has already been observed in its ancestors. AF2, for example, even predicted a novel fold (Durairaj et al. 2023), which is remarkable as experimental scientists discover fewer and fewer novel folds each year (Liisa Holm, personal communication). The Rost’s group introduced the notion of a dark proteome as the regions in protein space for which no structure information is available at all (Perdigão et al. 2015). Porta-Pardo *et al* recently showed that AF2 can illuminate the dark proteome, e.g. the fraction of dark proteome is reduced from 26% to just 10% when AF2 models are considered (Porta-Pardo et al. 2022). Experimental determination of many new amyloid structures, and training of AF2 and AF3 with these included into the training sets, could lead to further illumination of the dark amyloidome.

Ever since (Sander and Marks, 2012) and (Jones et al. 2012) showed that correlated mutations can be used for automatic structure prediction, a large part of the focus has been on ensuring that correlations really indicate direct, intra-molecular residue contacts. (Gouldson, Snell, and Reynolds 1997), for example, demonstrated that correlated mutations could be used to predict domain-swapped dimers for G Protein-Coupled Receptors. This work, though, involved a lot of manual intervention that cannot easily be automated, and it is just the type of exception that should be avoided when modelling proteins in general. AF2 and AF3 continued where Sander, Marks, Jones et al. stopped and fed the MSA - including the correlated mutations - to a structure predicting neural network. Information in the MSA about the amyloid fibril, if any, is likely to be completely overshadowed by signals from the globular fold. This problem was described for homooligomers predictions in (Uguzzoni et al. 2017). Larger proteins have, on average, more contacts per amino acid than smaller ones, and indeed AF3 more frequently predicted an amyloid structure for shorter proteins than for longer ones, probably because when fewer contacts are predicted, fewer signals are sent to AF3 that tell it to predict a globular fold.

AF3 is presently the best protein structure modelling software available. When predicting amyloid proteins with already known structures, AF3 outperformed Boltz and RibbonFold on CsgA and α-synuclein, though not on Aβ-42, for which RibbonFold performed best. For the rest of the well-known cases, none of the protocols managed to predict a fibrillar structure overlapping with the real one.

We think that a few actions can be taken to improve the AF3 predictions of amyloid proteins. These include using shorter sequences, predicting models of different oligomeric size, comparing results of the monomeric and oligomeric predictions, verifying if a homolog structure has been deposited in the PDB and whether it was used in AF3 training, running the prediction multiple times, careful interpretation of the model with metrics other than pLDDT, and testing different AF3 versions (e.g. desktop vs web sever). None of these, however, solve the problem of low data availability. Therefore, we can only hope that the experimental structure community will rapidly add many new amyloid fibril structures to the PDB. In the meantime, we suggest that amyloid structures are predicted in a three-step process: 1) first applying sequence analysis software to predict which fragments will form an amyloid fibril; 2) promote modeling of an amyloid form by defining the oligomeric state parameter, *n*; this parameter will provide some guidance, but additional human intervention through input specifications will likely be required in the future; 3) using a variety of structure prediction algorithms to model the relevant fibril in 3D and repeating the procedure a couple of times to generate multiple models.

It seems obvious that in the age of AI an automatic agent, trained on the scientific literature, could do a marvelous job in determining if amyloid structure prediction is needed at all or not. Maybe, screening for novel amyloid proteins will become routinely possible with AF4 or a higher version, as AF3 already occasionally predicts amyloid fibrils *de novo*. Still, amyloid model structures obtained with AF3 can already be useful in biomedical research, supporting hypothesis generation on amyloid protein structure-function relations, activity, or interactions with other molecules, thereby creating new insights into the role of amyloids in health and disease.

## Methods

### Datasets preparation

For the graphical summary of dataset preparation process, see Figure S16.

#### Positive-Control

To extract amyloid proteins without a solved experimental structure (the *Positive-control* dataset), three datasets, AmyPro (Varadi et al. 2018), the database of bacterial functional amyloids BFA (Nowakowska et al. 2023) and AmyLoad (Wozniak and Kotulska 2015), were used. AmyPro is an annotated database of amyloid proteins; it was last updated August 2023, according to their website. The BFA dataset was built based on the literature review in our previous work. In the BFA, no sequence apart from CsgA has a resolved structure. CsgA was excluded from this dataset. AmyLoad is a database of aggregating and non-aggregating sequences, many of which are short. However, no information about the PDB files for these sequences is provided. Given that there are few amyloid structures solved, even if some sequences from the AmyLoad have an amyloid structure in the PDB, they should constitute the minority.

All sequences of alpha phenol-soluble modulins were not considered, as these form amyloid fibrils with cross-alpha architecture (Tayeb-Fligelman et al. 2017). Seven proteins, namely Aap, Bap, Esp, PAc, SasG, YghJ, and Agglutinin-like protein 3, were too long for multimeric predictions with AF3 and hence were removed. Sequences with lengths below 10 amino acids were removed. This dataset was clustered at a 90% level of identity with cd-hit default parameters (Fu et al. 2012). This was performed to remove the bias towards certain proteins, which might be extensively studied with different mutations and fragments of their sequence.

Homologs of the *ResolvedAmyloidStructure* dataset (see below) in the *Positive-Control* dataset were identified with MMseqs2 easy-search with default parameters (minimum identity threshold of 30%) (Mirdita, Steinegger, and Söding 2019), and sequences appearing as hits were removed, making the final version of the *Positive-Control* dataset, including 153 sequences.

#### Negative-Control

The non-aggregating peptides (*Negative-control* dataset) were extracted from the AmyLoad. These sequences have been experimentally tested and showed no amyloid formation. As mentioned, it was impossible to identify if AmyLoad sequences have solved structures in the PDB.

Sequences with lengths below 10 amino acids were removed. Homologs of the *ResolvedAmyloidStructure* dataset in the *Positive-Control* dataset were identified with MMseqs2 easy-search with default parameters, and sequences appearing as hits were removed, making the final version of the *Negative-Control* dataset, including 56 sequences.

#### ResolvedAmyloidStructure and CsgA

The *ResolvedAmyloidStructure* dataset consisted of six proteins: Aβ-42, α-synuclein, glucagon, iapp (human amylin), transthyretin, and immunoglobulin. All these proteins have a structure solved for (almost) the entire sequence. The exact sequences are provided in Supplementary Table S2. The CsgA sequence is also given in Table S2.

### Structure prediction with AF3 web server

#### Positive-control and Negative-control

One AF3 structure with six protein copies (*copies -> 6*) and one monomer (*copies -> 1)* was predicted for each sequence from the *Positive-control and Negative-control* datasets. Template settings were set to *template settings* -> *Use PDB templates with default cut-off date (30/09/2021)* in the main run of the predictions. Additionally, the predictions were repeated with *template settings -> Turn off*. The *Entitity type* parameter was set to Protein. All models were predicted with the AF3 webserver available at https://alphafoldserver.com/welcome.

##### ResolvedAmyloidStructure and CsgA

Fifty AF3 models with six protein copies (*copies=6*) and fifty monomeric models (*copies=1)* for each protein in *ResolvedAmyloidStructure* were predicted. For Aβ-42, additionally, 50 structures per each number of protein copies from 1 to 9 were generated. Template settings were set to *template settings*=”*Use PDB templates with default cut-off date (30/09/2021)”.* The *Entitity type* parameter was set to Protein. All models were predicted with the AF3 webserver available at https://alphafoldserver.com/welcome.

### Structure prediction with other models

Other structure prediction algorithms than AF3 in the webserver version were used for six proteins in ResolvedAmyloidStructure and CsgA.

Five models per protein were generated with one ColabFold run of AF-Multimer (model_type=alphafold_multimer_v3) for each of these seven proteins (Mirdita et al. 2022, Evans et al. 2022) with three protocols: when the custom template was provided, without a template and with a template found in PDB100. The custom template used corresponds to the PDB identifiers provided in Table S2. For CsgA, the 8enr structure was used as a template. The rest of the parameters were used in a default mode. Ten models per protein were generated with the AF3 desktop version for these seven proteins; each model was produced with a different seed. The procedure was repeated with the same parameters, but with the custom MSA from ColabFold. The Multiple Sequence Alignments (MSAs) generated with ColabFold were used in the further predictions.

For transtyretin and immunoglobulin, the number of effective sequences in the MSA was calculated using the NEFFy software with default parameters (Haghani, Bhattacharya, and Murali 2025). The number of effective sequences reflects the diversity of sequences of an MSA. Four additional MSAs were created for these two proteins with a ‘full’ depth as well as 1000, 500 and 100 first sequences and for each version we calculated how many neffs there are (Table S3). For each of these MSA versions, ten structures were predicted for these two proteins with the desktop version of AF3 with default parameters.

Ten models per protein were generated with Boltz (Passaro et al. 2025; Wohlwend et al. 2025) version 1 with default parameters. For each protein, a multimer with an oligomeric state of 6 was modeled using MSA generated with ColabFold.

Ten models per protein were generated with AF3 desktop version for these seven proteins in two cases: the custom MSA generated with ColabFold was provided to the model and with default MSA. The rest of the parameters were set to default.

Ten models per protein were generated with one run of RibbonFold (Guo et al. 2025). The custom MSA generated with ColabFold was provided to the model. The rest of the parameters were default.

All models, apart from RibbonFold, were produced with the oligomeric state of six. RibbonFold assumes each model to have an oligomeric state of five.

### Quality of AF3 models

The quality of the models was assessed with pTM, ipTM and pLDDT metrics. The pTM score, which stands for predicted TM-score, represents the quality of a complex prediction. pTM value belongs to the [0,1) interval, with 0 meaning the poorest pTM. The ipTM score, which stands for the interface-predicted TM-score, represents the quality of relative subunits prediction. Analogous to pTM, ipTM takes values from [0,1) interval, where 0 refers to the worst ipTM score. pLDDT, per-residue predicted local distance difference test, is a measure of local confidence of the model. pLDDT takes values from 0 to 100. A pLDDT score above 90 is *ver*y *high,* and both backbone and side chains are likely to be correctly predicted. Values from 70 to 90 refer to a *confident* model, those from 50 to 70 are considered as *low* and below *50, very low (Jumper et al. 2021)*.

### TM-score

The similarity between structures was assessed with the TM-score that takes values from the interval (0,1]; the higher the TM-score, the more similar the structures. Structures with the TM-score>0.5 are assumed to be of the same fold in CATH (Xu and Zhang 2010).

### Structural clustering of the models

Models were clustered using Foldseek (version 09ce330d558c9479fe591a309676e74f519ed2f6) easy-cluster command with default parameters.

### Visualizations of models

Visualizations of all models were performed with PyMOL (version 3.0.3) and colored by the default AF colouring scheme, that is, according to the pLDDT (“PyMOL,” n.d.).

### Structure classification as amyloid

No reliable desktop tool is available for a large-scale assessment of whether the structure resembles an amyloid structure or not. Therefore for each model, we calculated its DSSP vector and the symmetry number. The dssp vectors were generated with Bio.PDB.DSSP module, available from Biopython (version 1.81), a DSSP object was defined with ‘mkdssp’ class (Cock et al. 2009). The symmetry groups were calculated with AnAnaS (Pagès and Grudinin 2020). The nomenclature of symmetry groups follows the AnAnaS paper. Models that were fully helical or showed c6 symmetry were considered nonfibrillar. For the remaining cases, manual curation was performed. A model was considered an amyloid structure when it was composed predominantly of beta-strands aligned perpendicularly to the axis of a fibril.

### Similarity of monomers, multimers, PDBs and STAMP

The PDB database was downloaded with Foldseek (van Kempen et al. 2024) in March, 2025 (Foldseek PDB version: df8de2b01408fe438e3f075698032906). Some of the entries in this PDB version have not been seen by AF3. However, they constitute only a minority of the database. Models were compared to amyloid structures available in STAMP database (Louros et al. 2022). STAMP is a manually-curated database of amyloid structures available in the PDB. STAMP does not contain recently solved structures of CsgA - 8enr and 8enq. Therefore, in our analyses, we extended STAMP by these two PDB files.

Foldseek compares structures to detect even distant homology relationships. It has been used with default parameters throughout this study. Foldseek easy-search returns structure file identifiers and a description of the structural alignment between the input structure and the 3D aligned part of that PDB file. Foldseek hits can be described by TM-score and homolog probability, which stands for “the probability for each match to be homologous, based on a fit of true and false matches on SCOPe” (direct quote from van Kempen, 2024). When mentioned, this list of hits was filtered to include only 3D alignments with TM-score>0.5. Hits discovered by Foldseek on AF2 models often relate to hits found by traditional sequence-based approaches like BLAST (Monzon et al. 2022).

#### Positive-control and Negative-control

Each monomer from the *Positive-control and Negative-control* datasets was compared with its corresponding multimer with Foldseek easy-search with default parameters (*easy-search protein_monomer protein_multimer output tmp*). Similar structures to *Positive-control* multimeric models were searched in the PDB with Foldseek easy-search with default parameters (*easy-search control_multimers pdb output tmp*). Similar sequences to *Positive-control* were searched in the PDB with MMseqs2 (version 13-45111+ds-2) easy-search with default parameters (*easy-search control_multimers pdb output tmp*) (Mirdita, Steinegger, and Söding 2019).

#### ResolvedAmyloidStructure

Fifty monomeric structures predicted for each protein from the *ResolvedAmyloidStructure* were clustered with Foldseek easy-cluster with default parameters. For each representative monomeric structure, for each protein, similar structures were searched among fifty corresponding multimeric predictions with Foldseek easy-search with default parameters (*easy-search representative_protein_monomer protein_multimers output tmp*). Similar structures to *ResolvedAmyloidStructure* multimeric models were searched in the PDB with Foldseek easy-search with default parameters (*easy-search ResolvedAmyloidStructure_multimers pdb output tmp*). Similar structures to *ResolvedAmyloidStructure* multimeric models were searched in STAMP with Foldseek easy-search with default parameters (*easy-search ResolvedAmyloidStructure_multimers stamp output tmp*). Hits regarding only the same protein were considered in the analyses.

##### Visualizations of structure alignments

Visualization of structure alignments was performed with the Foldseek Search Server https://search.foldseek.com/search.

### General data analysis

All data analyses were performed in Python 3.11 using the following packages: pandas version 2.2.3 (Reback *et al*., 2020), biopandas version 0.4.1 (Raschka, 2017), numpy version 1.26.4 (Harris et al. 2020), matplotlib version 3.10.0 (Hunter & Dale, 2007), seaborn version 0.13.2 (Waskom, 2021), scikit-learn version 1.6.1 (Kramer, 2016) and scipy version 1.15.1 (Virtanen *et al*, 2021).

## Data availability

All predictions for the *Positive-control, Negative-control*, *ResolvedAmyloidStructure* datasets and CsgA, along with the metadata, and MSA files are available at the Zenodo repository: https://zenodo.org/records/17544325. All bash one-liners, scripts, and other software written to perform this study are available upon request from AWW.

## Conflict of interest statement

The authors declare no conflicts of interest.

## Funding

This work was supported by the National Science Centre, Poland [2019/35/B/NZ2/03997].

## Acknowledgements

We gratefully acknowledge Polish high-performance computing infrastructure PLGrid (HPC Center: ACK Cyfronet AGH) for providing computer facilities and support within computational grant no. PLG/2025/018263. MK would like to acknowledge the fellowship from Stellenbosch Institute for Advanced Study (STIAS).

## Notes

### Competing Interest Statement

The authors have declared no competing interest.

### Summary of Updates

The manuscript has been revised and improved to a VERY LARGE EXTENT.

