## Supplementary for "Non-standard proteins in the lens of AlphaFold 3 - a case study of amyloids"

### CONTENT

**Figure S1.** Distribution of the oligomeric states of 577 amyloid structures in STAMP.
**Figure S2.** Distributions of AF3 quality metrics for all datasets.
**Figure S3.** A case study of Alpha-Spectrin SH3 domain deposited in AmyLoad.
**Figure S4.** Distribution of the sequence length for *correct* and *wrong* AF3 predictions for the *Positive-control*.
**Figure S5.** Confusion matrix for the predictions of the *Positive-control* and *Negative-control* datasets.
**Figure S6.** Sequence length distribution and similarity between monomeric and multimeric models for the *Positive-control* dataset.
**Figure S7.** AF3 models for amyloids match PDB structures, harnessing fibrillar structure prediction.
**Figure S8.** Cluster representatives for *ResolvedAmyloidStructure* and CsgA proteins.
**Figure S9.** Best Foldseek hits for each AF3 model for *ResolvedAmyloidStructure* dataset and CsgA in the STAMP dataset of amyloid structures.
**Figure S10.** A list of the best Foldseek hits for AF3 models in the PDB.
**Figure S11.** The cluster representative of all AF3 models for transthyretin and immunoglobulin.
**Figure S12.** AF-Multimer models for the *ResolvedAmyloidStructure* dataset (and CsgA).
**Figure S13.** ipTM score distribution for *ResolvedAmyloidStructure* and CsgA models produced with AF-Multimer.
**Figure S14.** pLDDT score distribution for *ResolvedAmyloidStructure* and CsgA models produced with AF-Multimer.
**Figure S15.** Comparison of AF3 performance to other structure prediction algorithms for Aβ-42.
**Figure S16.** Summary of the dataset preparation process.

**Table S1.** Mmseqs hits found for the sequences of correct *Positive-control* models in the PDB.
**Table S2.** Sequences of *ResolvedAmyloidStructure* datasets and examples of PDB identifiers of amyloid structures that were part of the AF3 training set.
**Table S3.** Number of effective sequences (Neff) in the Colab-generated MSA and in the cases when only the first 1000, 500 and 100 sequences of it were extracted and applied in AF3 modelling.

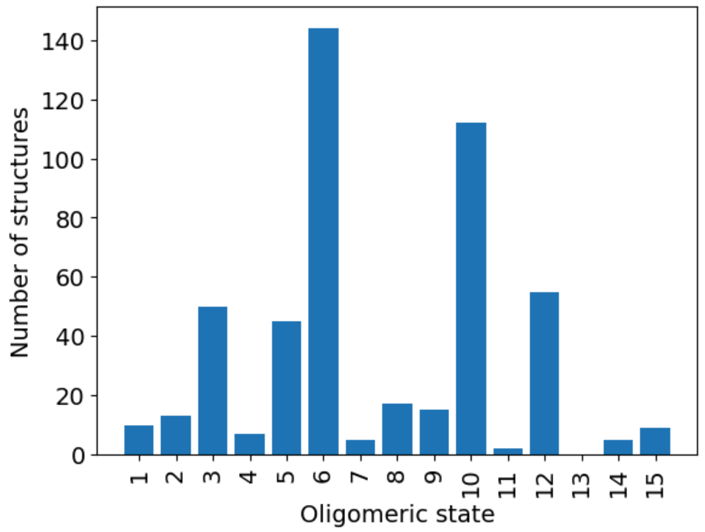


**Figure S1. Distribution of the oligomeric states of 577 amyloid structures in STAMP (database of amyloid structures in the PDB).** 88 more PDB entries with an oligomeric state (much) larger than 15 were not included.

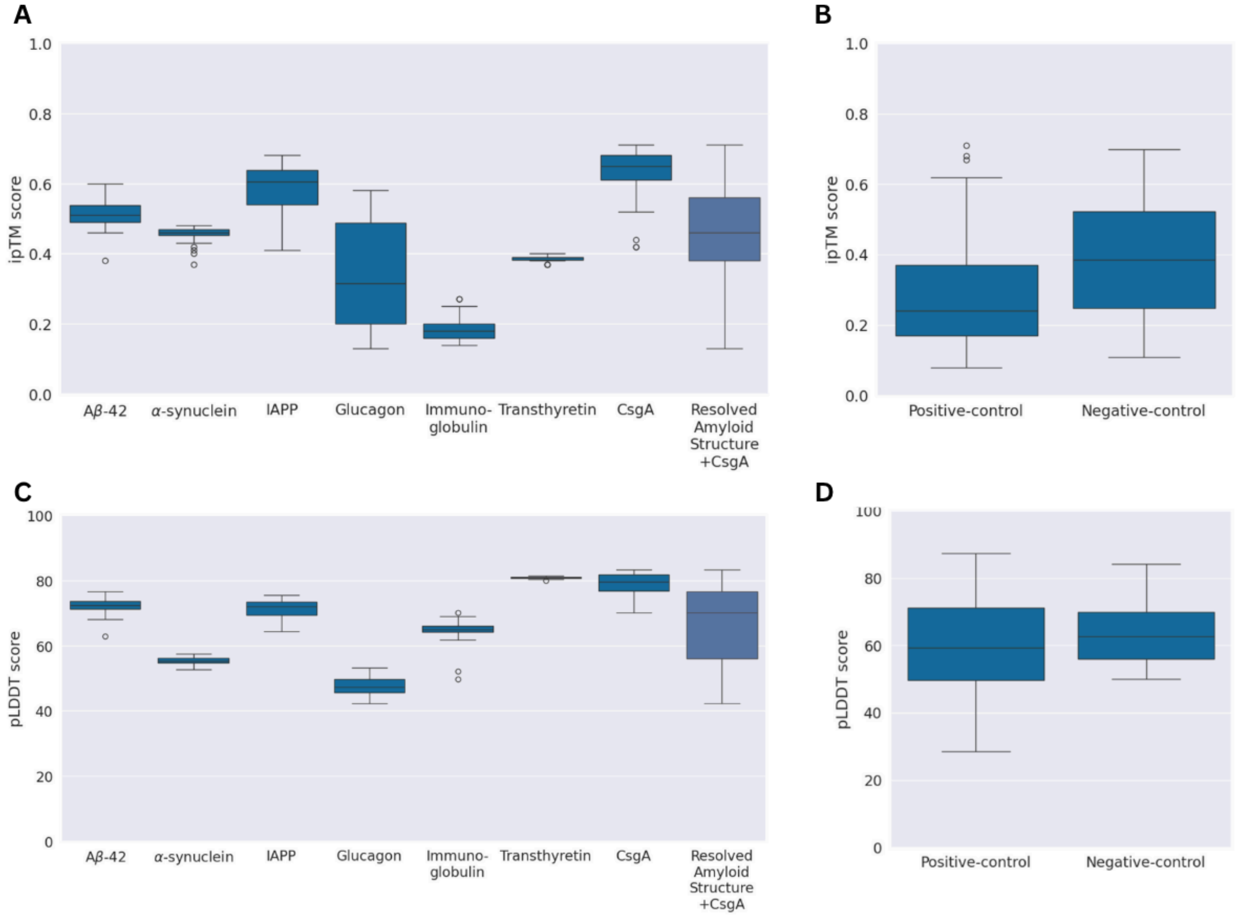


**Figure S2. Distributions of AF3 quality metrics for all datasets.** A. ipTM score distribution for *ResolvedAmyloidStructure* and CsgA. B. ipTM score distribution for *Positive-* and *Negative-control*. C. pLDDT score distribution for *ResolvedAmyloidStructure* and CsgA. D. pLDDT score distribution for *Positive-* and *Negative-control*.

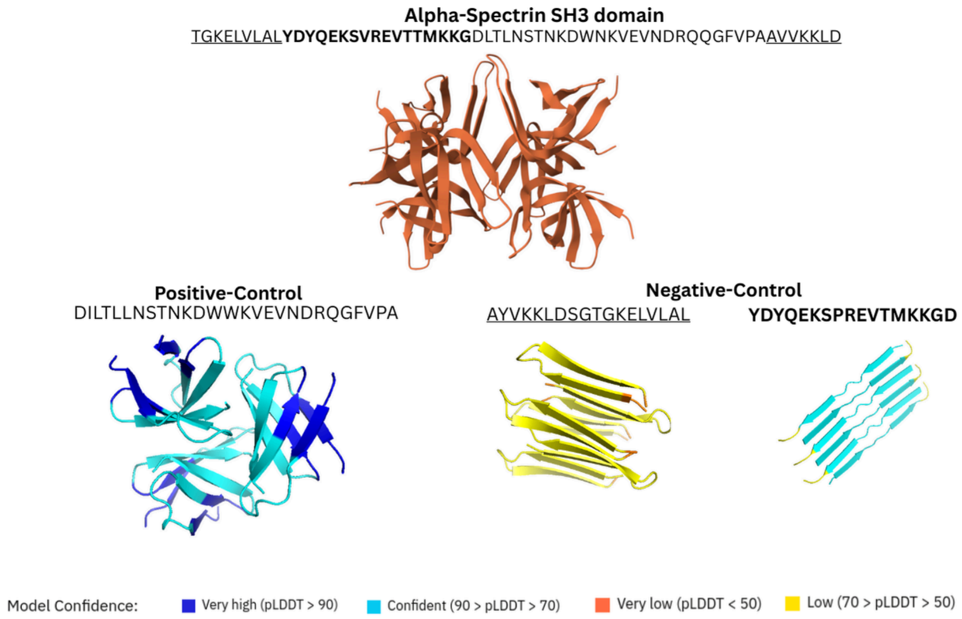


**Figure S3. A case study of Alpha-Spectrin SH3 domain deposited in AmyLoad.** The entire structure of the domain is not predicted as an amyloid. The aggregation-prone region (present in *Positive-control* dataset) is not predicted as an amyloid. The non-aggregating peptides of this domain (present in *Negative-control* dataset) are both predicted as amyloid structures.

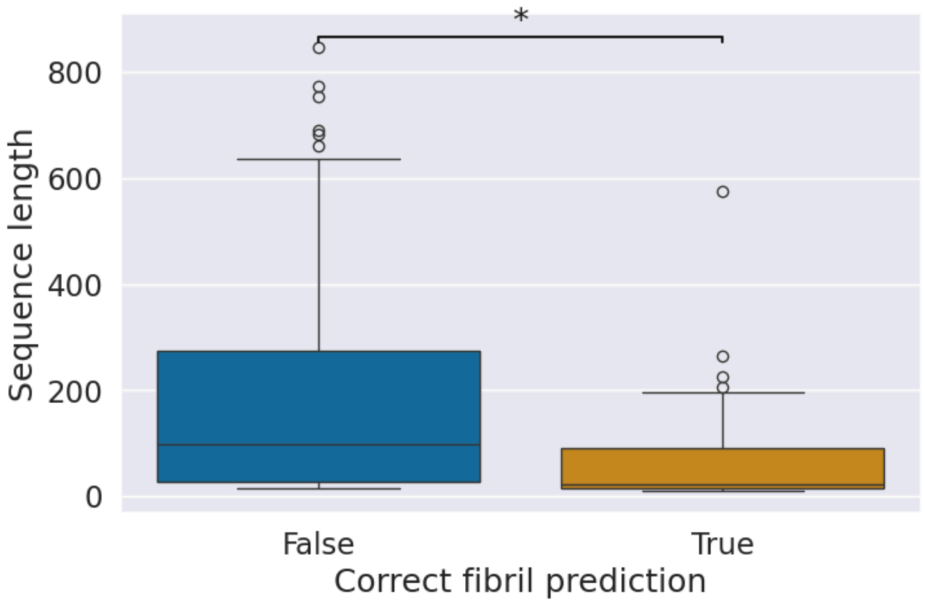


**Figure S4. Distribution of the sequence length for *correct* and *wrong* AF3 predictions for the *Positive-control*.**

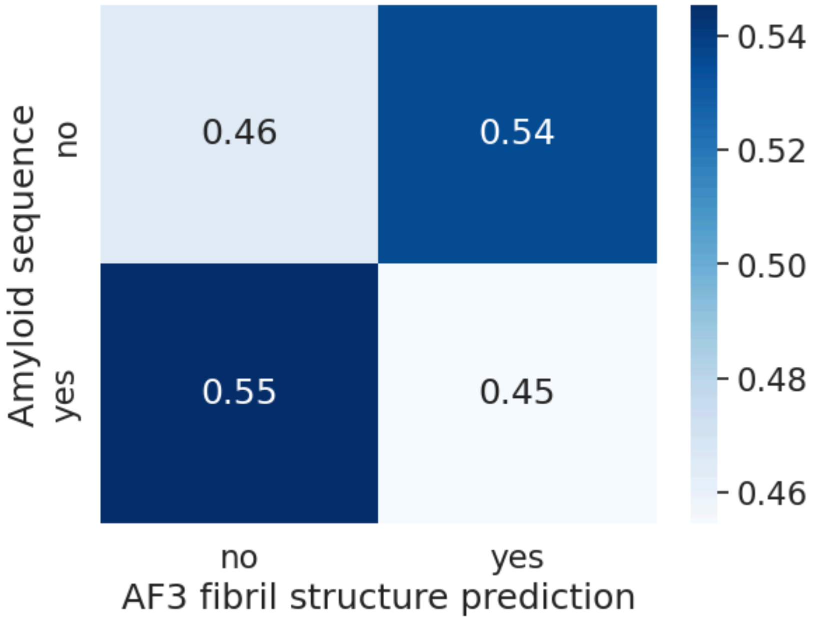


**Figure S5. Confusion matrix for the predictions of the *Positive-control* and *Negative-control* datasets.** To create this confusion matrix, we used only sequences of lengths 36 or shorter in the *Positive-control*. All sequences in the *Negative-control* are also of length 36 or shorter.

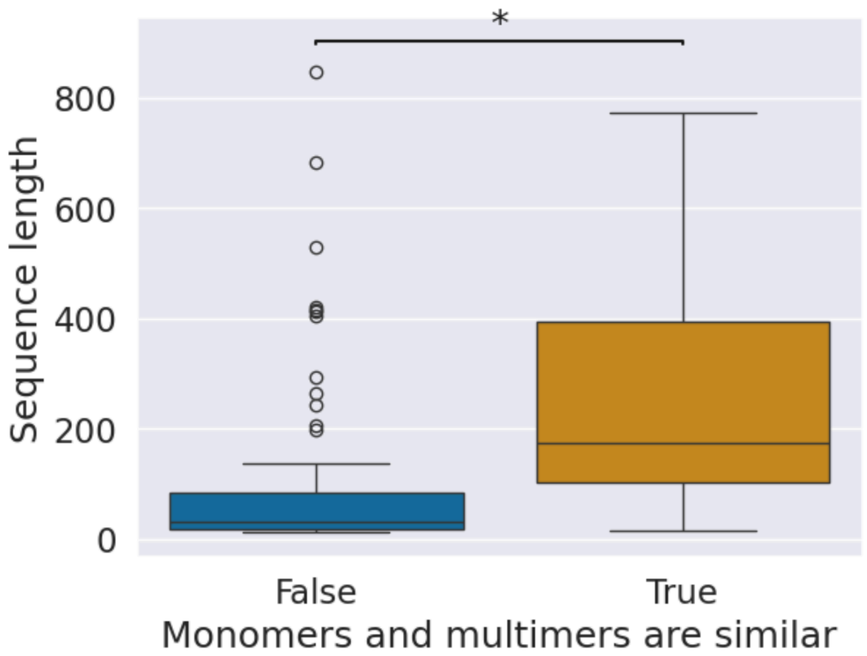


**Figure S6. Sequence length distribution and similarity between monomeric and multimeric models for the *Positive-control* dataset.** The difference between the left and right distributions was statistically significant; Mann-Whitney p-value=3e-10. When models for sequences with a length above 36 amino acids are only considered, the similarity between multimeric and monomeric models implies an increase in the pLDDT score by 25 points (statistically significant, Mann-Whitney p-value=2e-12).

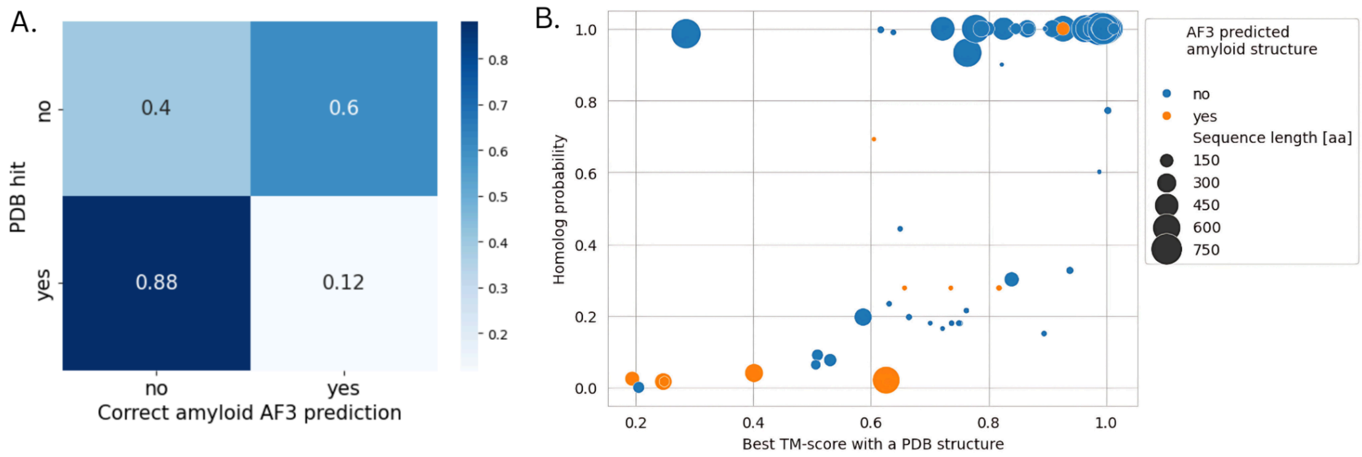


**Figure S7. AF3 models for amyloids match PDB structures, harnessing fibrillar structure prediction. A.** Confusion matrix for amyloid proteins from the *Positive-control* dataset concerning AF3 predictions and similarity of the models to PDB structures. **B.** Results of the search of AF3 models in the PDB for amyloid proteins (Positive-control). Foldseek homolog probability as a function of the TM-score between the AF3 model and the most similar PDB file. Higher homolog probability was associated with higher pLDDT (mean pLDDT with a homolog probability above 0.95 was 67, with no homolog 57, p-value=1e-4).

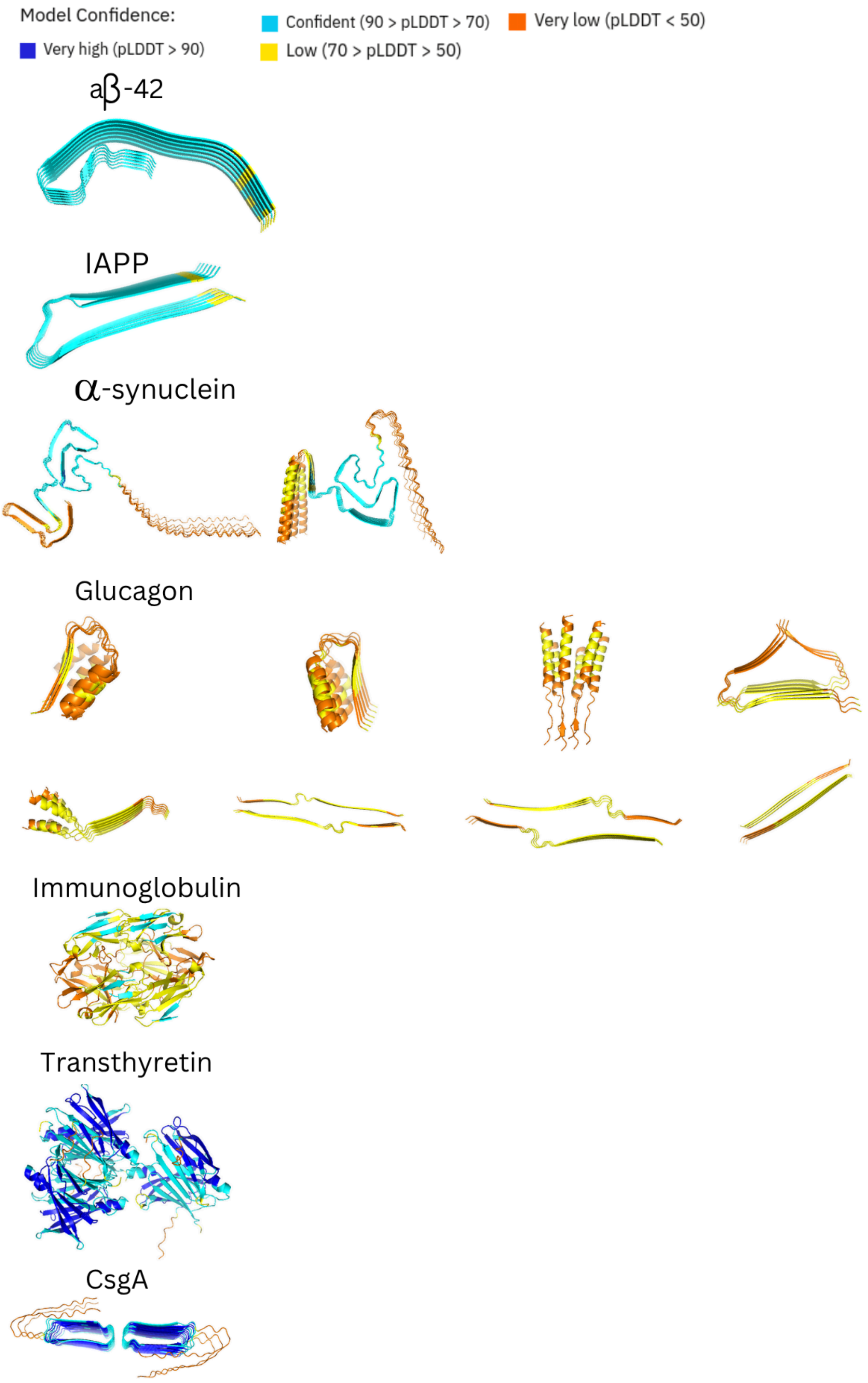


**Figure S8. Cluster representatives for *ResolvedAmyloidStructure* and CsgA proteins.** Structures are coloured according to the pLDDT score per residue.

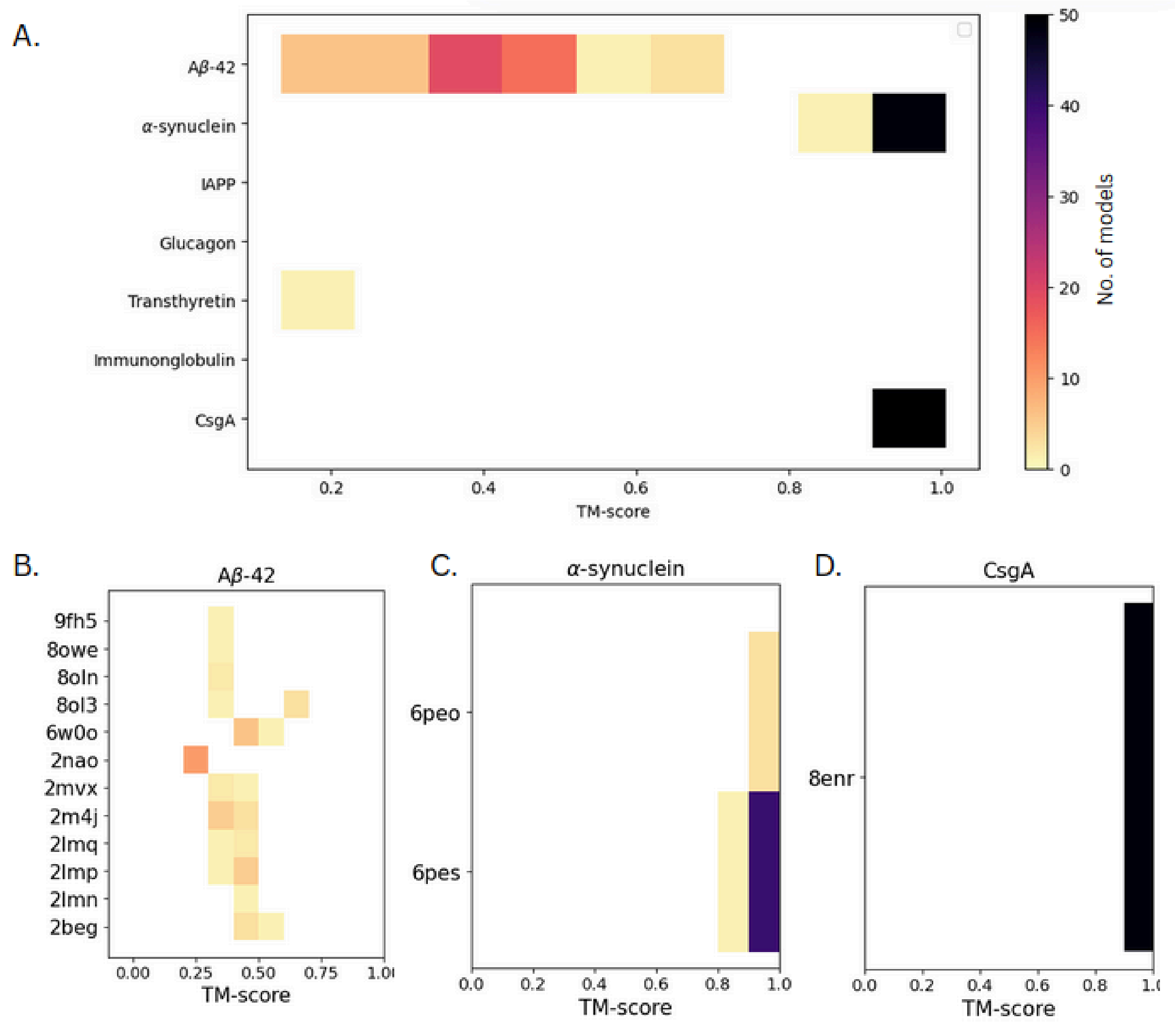


**Figure S9. Best Foldseek hits for each AF3 model for *ResolvedAmyloidStructure* dataset and CsgA in the STAMP database of amyloid structures.** The colors represent the number of models that had the best match to a PDB identifier. **A.** TM-scores of all hits. **B.** Similarity between 50 Aβ-42 AF3 models and amyloid structures of this protein in STAMP. **C.** Similarity between 50 α-synuclein AF3 models and amyloid structures of this protein in STAMP. **D.** Similarity between 50 CsgA AF3 models and amyloid structures of this protein in STAMP (extended by 8enr and 8enq PDB files).

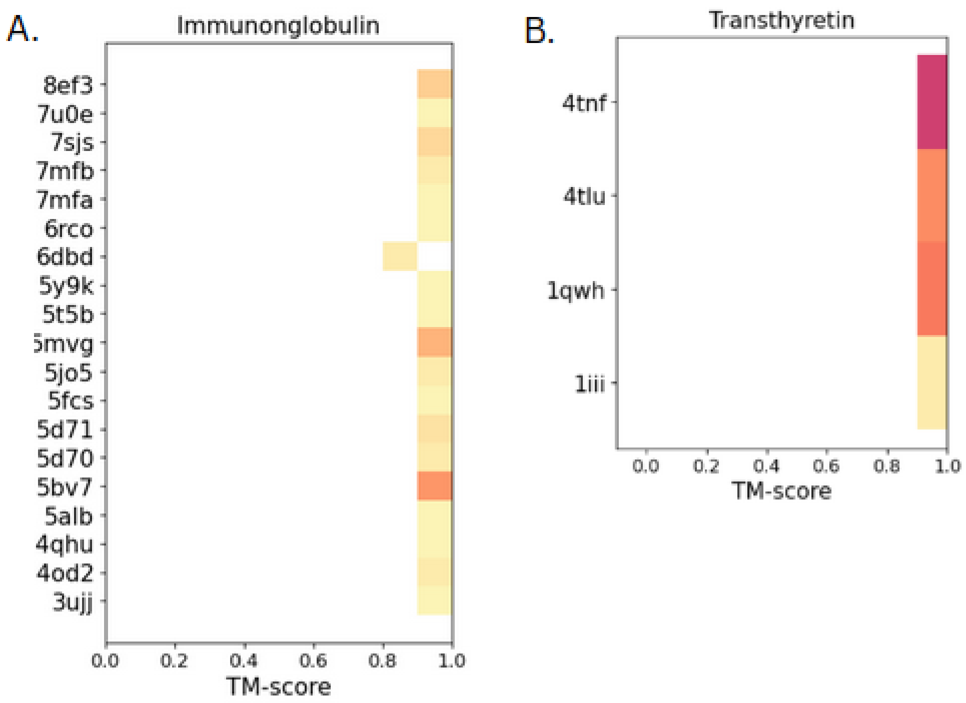


**Figure S10. A list of the best Foldseek hits for AF3 models in the PDB** for A. immunoglobulin and B. transthyretin. All hits are globular structures.

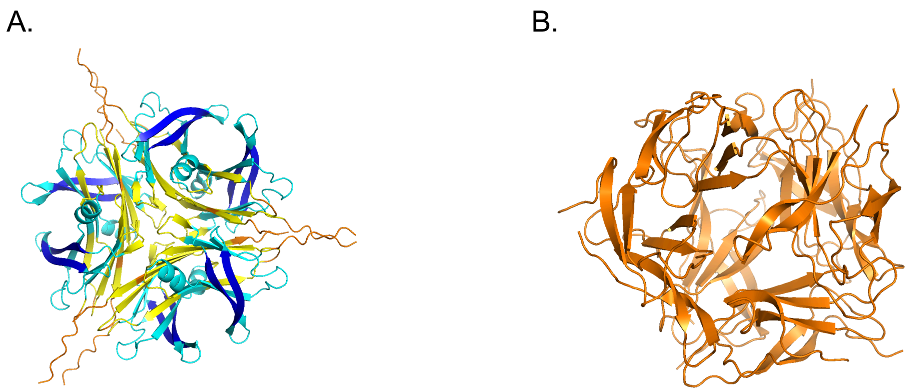


**Figure S11. The cluster representative of all AF3 models** for A. transthyretin and B. immunoglobulin.

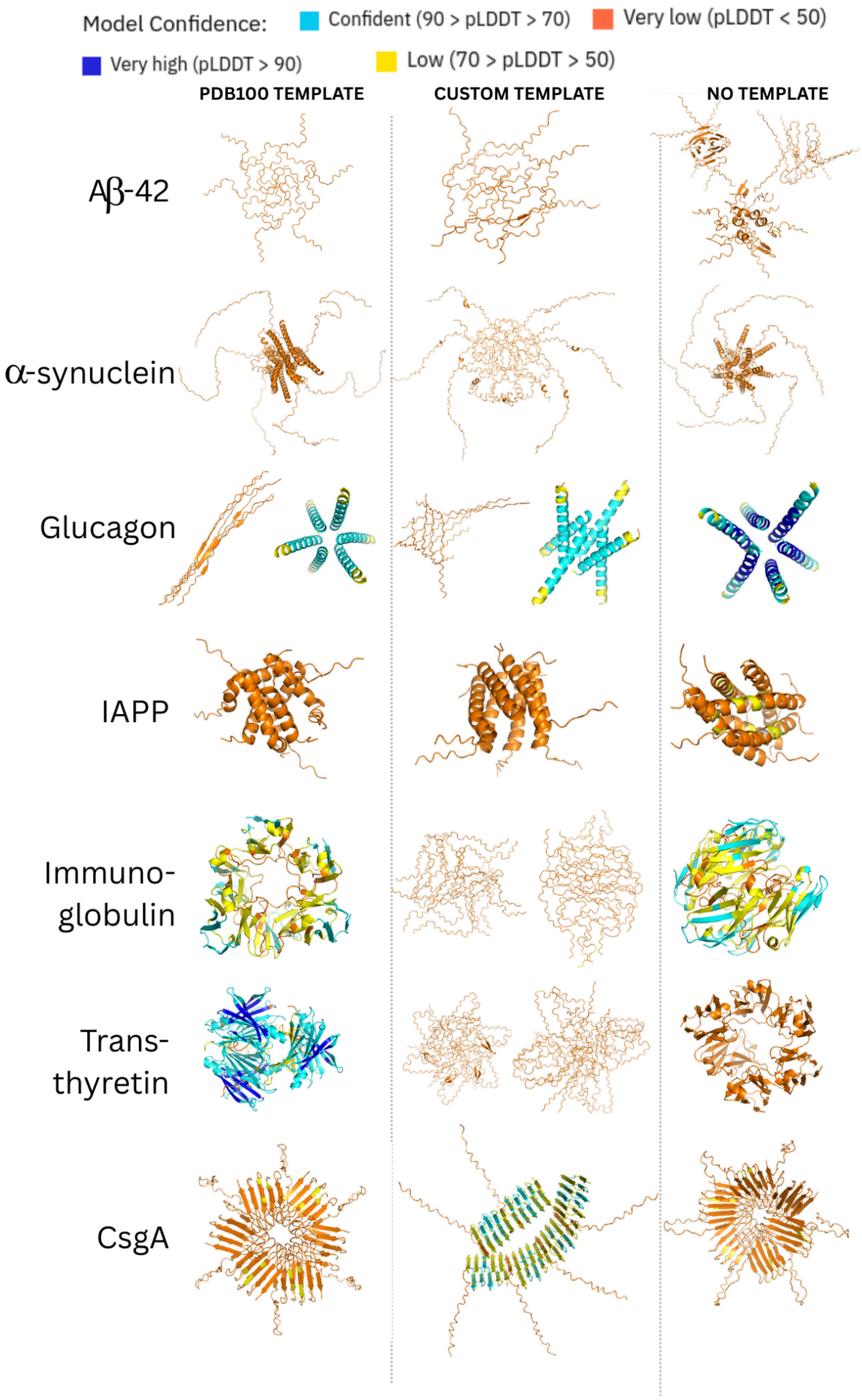


**Figure S12. AF-Multimer models for the *ResolvedAmyloidStructure* dataset (and CsgA):** using a PDB100 template, a custom template from Table S2 and without any template**.** Five models generated with AF-Multimer in each case were clustered with Foldseek and their cluster representatives are shown.

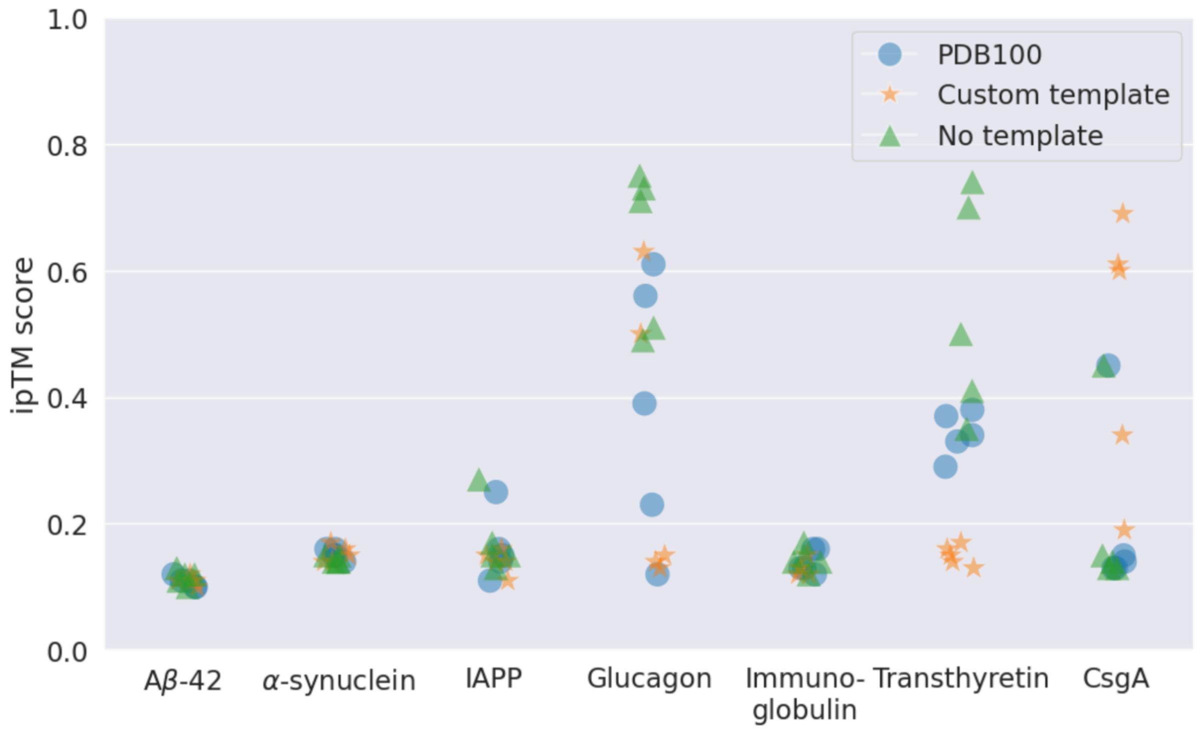


**Figure S13. ipTM score distribution for *ResolvedAmyloidStructure* and CsgA models produced with AF-Multimer.**

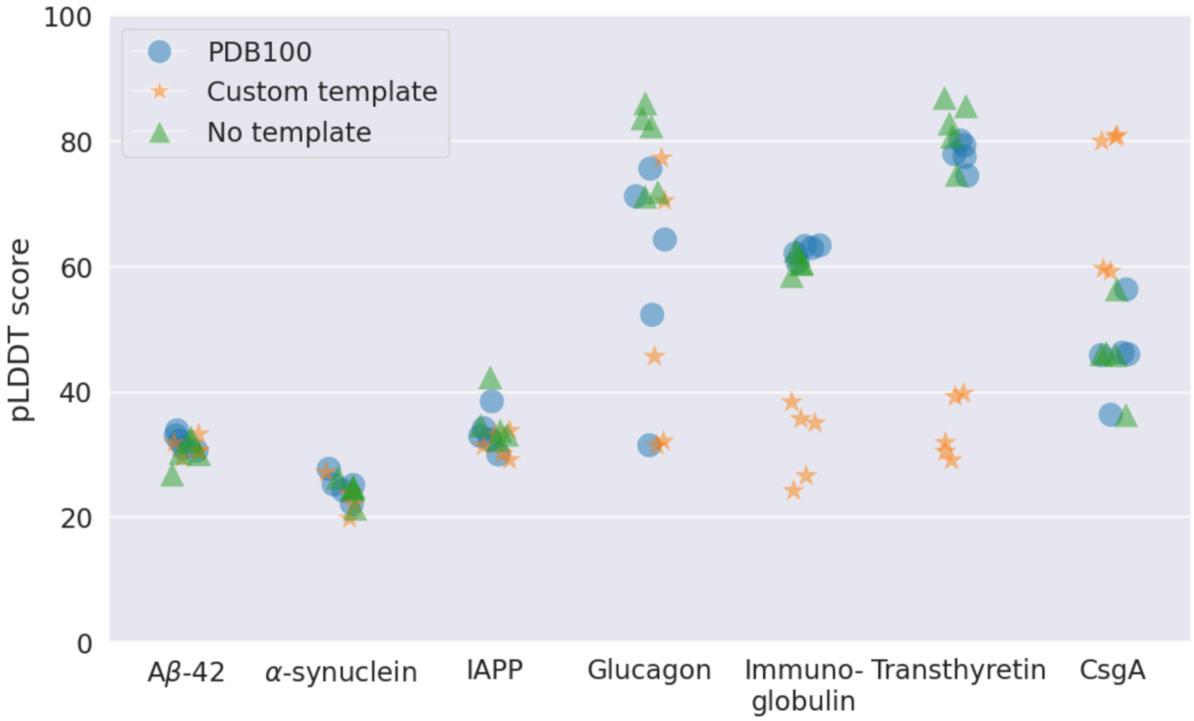


**Figure S14. pLDDT score distribution for *ResolvedAmyloidStructure* and CsgA models produced with AF-Multimer.**

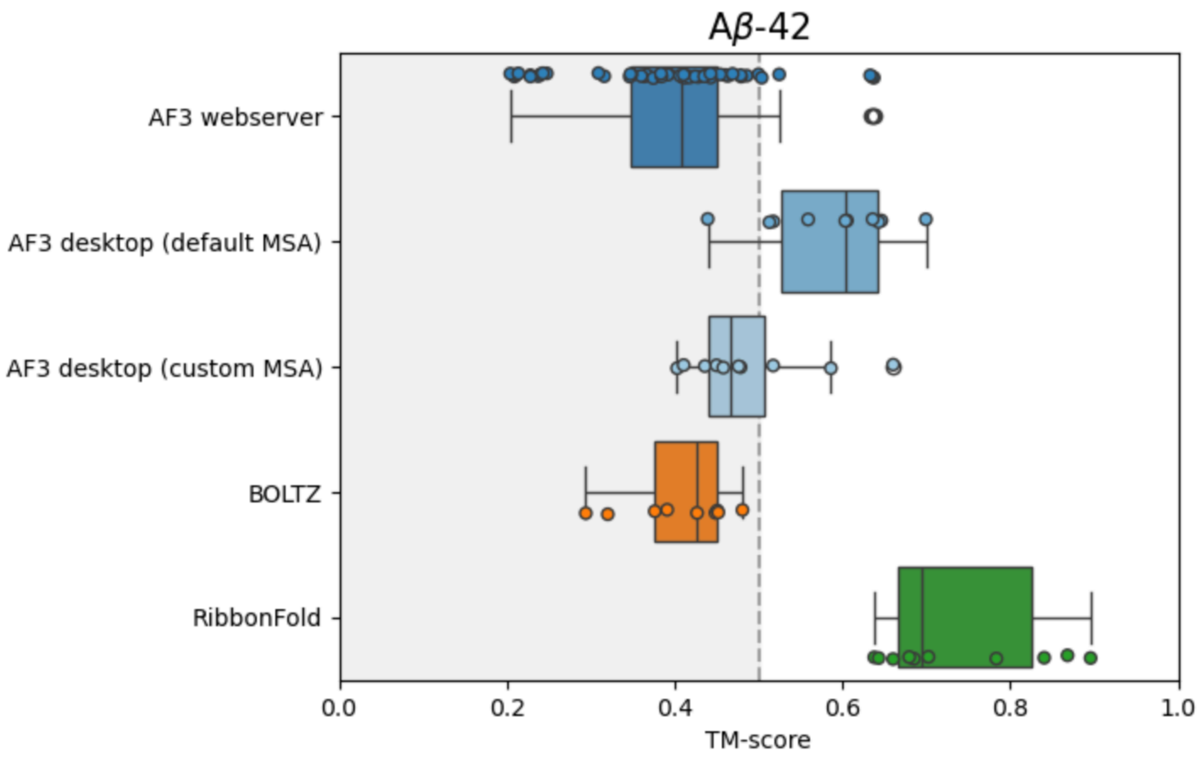


**Figure S15. Comparison of AF3 performance to other structure prediction algorithms for Aβ-42.** Models were aligned with Foldseek to the solved amyloid structures of Aβ-42 available in STAMP. TM-score of the best hit for each model is provided.

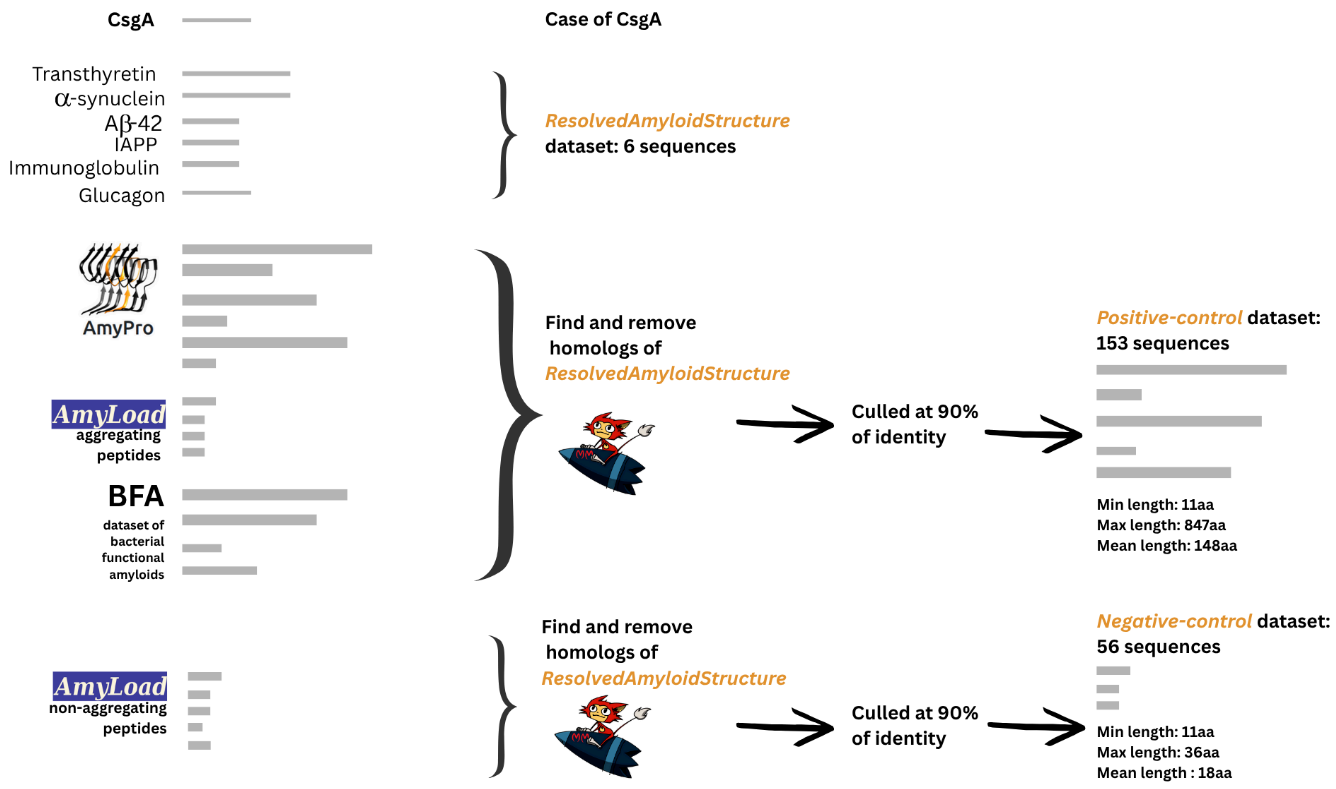


**Figure S16. Summary of the dataset preparation process.**

| **Protein** | **AF3 multimer model (models colored by pLDDT)** | **mmseqs hit in the PDB** | **mmseqs alignment E-value, (identity)** | **Comment about the hit PDB file** |
|---|---|---|---|---|
| Microcin E492 | 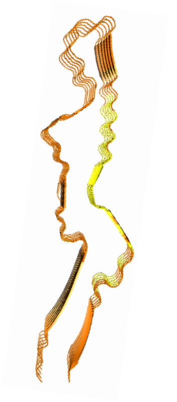 | 7dyr 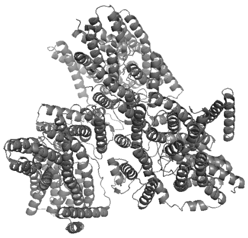 | 1.7e-42 (1.0) | Protein complex with globular microcin |
| PB1-F2, C-terminal domain | 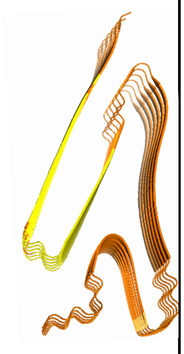 | 2hn8 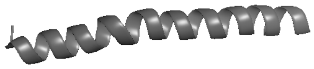 | 4e-19 (0.95) | Globular monomer of the protein with the C-terminal fragment |
| Defensin-like protein 1, C-terminal peptide | 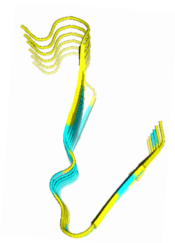 | 2n2r 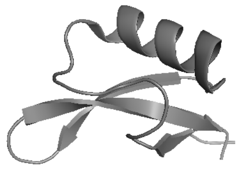 | 2.3e-17 (0.97) | Globular monomer of a homolog |
| Obestatin | 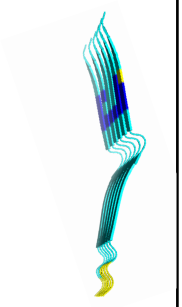 | 2jsj 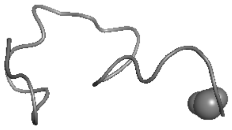 | 6.8e-8 (0.87) | Globular monomer of a homolog |

| | | | | |
|---|---|---|---|---|
| Merozoite surface protein 2, MSP2 | 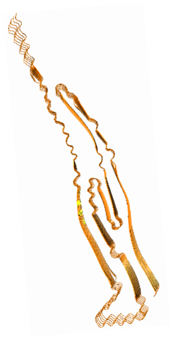 | 2mu8 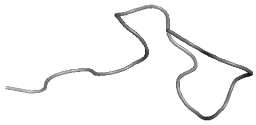 | 6.3e-4 (1.0) | Globular monomer of a small fragment of this protein |
| Calcitonin | 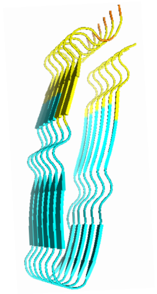 | 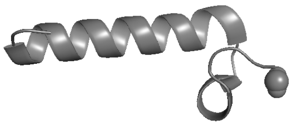 | 4.9e-14 (0.85) | Non-amyloidogenic analogue of the protein |
| Alpha-S2-casein | 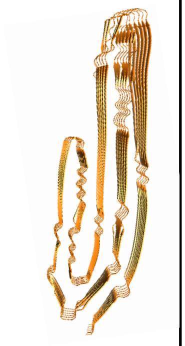 | 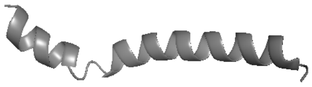 | 5.5e-15 (1.0) | Distant homologous globular fragment of a protein |
| Natriuretic peptides B, ProBNP | 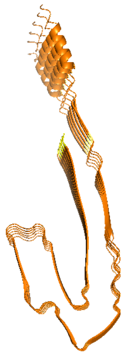 | 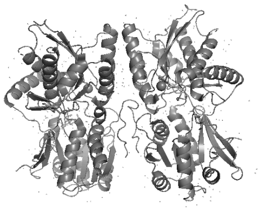 | 1.4e-6 (1.0) | Homologous globular fragment of a protein |
| Medin | 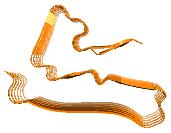 | 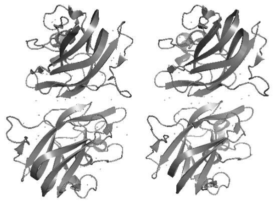 | 2.4e-17 (0.66) | Globular monomer of a homolog |

**Table S1. Mmseqs hits found for the sequences of correct *Positive-control* models in the PDB.**

| Protein | Sequence | Example of a PDB identifier of an amyloid structure that was part of the AF3 training set |
|---|---|---|
| Aβ-42 | DAEFRHDSGYEVHHQKLVFFAEDVGS NKGAIIGLMVGGVVIA | 2mxu |
| α-synuclein | MDVFMKGLSKAKEGVVAAAEKTKQGV AEAAGKTKEGVLYVGSKTKEGVVHGV ATVAEKTKEQVTNVGGAVVTGVTAVA QKTVEGAGSIAAATGFVKKDQLGKNE EGAPQEGILEDMPVDPDNEAYEMPSE EGYQDYEPEA | 2n0a |
| IAPP | KCNTATCATQRLANFLVHSSNNFGAIL SSTNVGSNTY | 6vw2 |
| Glucagon | HSQGTFTSDYSKYLDSRRAQDFVQW LMNT | 6nzn |
| Immunoglobulin lambda variable 3-19 light chain | AVSVALGQTVRITCQGDSLRSYSASW YQQKPGQAPVLVIFRRFSGSSSGNTA SLTITGAQAEDEADYYCNSRDSSANH QVFGGGTKLTV | 6z1i |
| Transthyretin | GPTGTGESKCPLMVKVLDAVRGSPAI NVAMHVFRKAADDTWEPFASGKTSES GELHGLTTEEEFVEGIYKVEIDTKSYW KALGISPFHEHAEVVFTANDSGPRRYT IAALLSPYSYSTTAVVTNPKE | 6sdz |
| CsgA | MGVVPQYGGGGNHGGGGNNSGPNS ELNIYQYGGGNSALALQTDCRNSDLTI TQHGGGNGADVGQGSDDSSIDLTQR GFGNSATLDQWNGKNSEMTVKQFGG GNGAAVDQTASNSSVNVTQCGFGNN ATAHQY | |

**Table S2. Sequences of *ResolvedAmyloidStructure* datasets and examples of PDB identifiers of amyloid structures that were part of the AF3 training set**. In the grey colour, the used sequence of CsgA is given.

| Protein | Depth of the full Colab MSA | Neff: Colab MSA full | Neff: MSA with first 1000 sequences | Neff: MSA with first 500 sequences | Neff: MSA with first 100 sequences |
|---|---|---|---|---|---|
| Immunoglobulin lambda variable 3-19 light chain | 13297 | 885 | 27 | 6 | 2 |
| Transthyretin | 6176 | 385 | 47 | 13 | 2 |

**Table S3. Number of effective sequences (Neff) in the Colab-generated MSA and in the cases when only the first 1000, 500 and 100 sequences of it were extracted and applied in AF3 modelling.**
